# Scalable Human Cellular Models of Parkinson’s Disease Reveal A Druggable Link Between the Angiotensin Receptor 1 and α-Synuclein Pathology

**DOI:** 10.1101/2025.11.18.689137

**Authors:** Mathieu Daynac, Vincent Mouilleau, Yuxi Liu, Xiaoyu Yang, Yin Shen, Su Guo

## Abstract

**Background:** Parkinson’s disease (PD) involves progressive loss of midbrain dopaminergic (mDA) neurons in the substantia nigra. No disease-modifying treatments exist, only symptomatic relief. Our lab reported an unbiased screen in larval zebrafish identifying renin-angiotensin-aldosterone system (RAAS) inhibitors, including clinically used AGTR1 inhibitors for hypertension, as potent neuroprotective agents. This study aims to investigate the effects of AGTR1 inhibition on human mDA neuron survival using inducible neurodegenerative 2D and 3D models for human mDA neuron degeneration.

**Methods:** We report a scalable high-content platform, using CRISPR-engineered human induced pluripotent stem cell (hiPSC)-derived mDA neurons expressing a tyrosine hydroxylase (TH) fluorescent reporter, allowing to track mDA neuron survival live “in a dish”. We developed chemically inducible neurodegenerative 2D and 3D models for human mDA neuron degeneration, allowing to recapitulate PD pathology in human cells *in vitro*.

**Results:** Our model establishes scalable human cellular models of PD well-suited for therapeutic discovery. Using 2D and 3D mono and co-cultures, this study demonstrates that inhibition of AGTR1, via chemical or genetic means, protects against chemically induced mDA neuron degeneration. Transcriptomic analyses show AGTR1 inhibition lowers synuclein transcription, by reducing *SNCA* and *SNCB* gene expression. In 3D neuron-glia assembloids, AGTR1 inhibition protects against the accumulation of phosphorylated form of α-synuclein (p129α-Syn), key PD pathological marker.

**Conclusions:** We highlight AGTR1 as a key regulator of α-synuclein transcription and aggregation in human mDA neurons, and AGTR1 inhibition as pro-survival in human iPSC-derived models. These findings position inhibition of AGTR1 as a promising therapeutic strategy for PD neuroprotection.

## BACKGROUND

Parkinson’s disease (PD) is a prevalent and progressive neurodegenerative disorder marked by the selective loss of dopaminergic (DA) neurons in the substantia nigra pars compacta (SNpc). This neuronal loss leads to debilitating motor deficits, including bradykinesia, rigidity, and resting tremor. Although current therapeutic strategies can alleviate these symptoms by enhancing dopaminergic signaling, their long-term efficacy is limited. Furthermore, they do not address the underlying process of neurodegeneration. Thus, there remains an urgent need for disease-modifying treatments that can halt or reverse dopaminergic neuron loss.

We developed a chemogenetic zebrafish model of DA neuron degeneration optimized for high-content screening and performed a screen of over 1,400 bioactive small molecules. This screen identified renin-angiotensin-aldosterone system (RAAS) inhibitors, including clinically approved Angiotensin Receptor 1 (AGTR1) inhibitors known as sartans, as robust neuroprotective agents [1,2]. The RAAS pathway is a peptide ligand-modulated G-protein coupled receptor (GPCR) system conserved across vertebrates and extensively studied in peripheral tissues in the context of blood pressure modulation and salt homeostasis [3]. The existence of brain RAAS [4] is further supported by recent single-cell transcriptomic analyses revealing AGTR1 expression in subtypes of human SNpc DA neurons that are most vulnerable to degeneration in PD patients [5].

In rodents, it has been shown that AGTR1 inhibition offers neuroprotection, particularly in DA neurons, by mitigating oxidative stress, neuroinflammation, and dopaminergic degeneration [6–9]. Conversely, overactivation of AGTR1 enhances oxidative stress and neuroinflammation, contributing to dopaminergic neuron degeneration [10–14].

Despite these advances in animal systems, whether AGTR1 inhibition exerts neuroprotection on human midbrain DA neurons is unknown. To address this, we established a high-content platform by differentiating hiPSCs into mDA neurons. We applied it to the tyrosine hydroxylase (TH)-TdTomato (TdTom) reporter-expressing KOLF2.1J line [15,16] to enable live-cell imaging and automated quantification in both 2D monolayers and 3D neuron-glia assembloid formats. We further developed robust neurodegeneration models induced by conduritol B-epoxide (CBE), an inhibitor of glucocerebrosidase (GBA1), the most common genetic risk factor for PD [17], and Rotenone (Rot), a widely utilized neurotoxin that triggers severe neuronal loss via mitochondrial complex I inhibition [18]. Our findings demonstrate that AGTR1 inhibition, with antagonists including candesartan and valsartan or CRISPRi of AGTR1, conferred significant protection against CBE- and rotenone-induced neurotoxicity in human mDA neuron models. Transcriptomic profiling revealed that AGTR1 inhibition downregulates *SNCA* and *SNCAIP*, genes encoding α-synuclein and its aggregation partner synphilin-1, as well as *SNCB*, providing a mechanistic link between RAAS signaling and a defining molecular feature of PD pathology.

## METHODS

### Cell lines

Human induced pluripotent stem cells (hiPSCs); THTdTom-CRISPR Knock-In hiPSC lines [16]; KOLF2.1J [15] were grown onto Growth Factor Reduced (GFR) Matrigel, phenol red free Matrigel (CORNING #356231) coated dishes with StemFlex™ Medium (Life Technologies #A33493-01). hiPSCs were grown and passaged every 4-5 days using accutase (GIBCO, #A11105-01). They were tested for potential mycoplasma contamination every other week (MycoAlertTM Mycoplasma Detection Kit, Lonza, LT07-118). hiPSCs were thawed in presence of the ROCK inhibitor Y-27632 (10 μM, Stem Cell Technologies, #72304) and the culture medium was changed every day.

### hiPSC diCerentiation into midbrain dopamine (mDA) neurons

After reaching 60-70% confluence, hiPSCs were dissociated into single cells with Accutase (GIBCO, #A11105-01) for 3-5 min at 37°C and resuspended in day 0 differentiation medium seeded in ultra-low attachment 6-well plates (Fisher Scientific, #3471) (4×10^5^ cells ml^−1^) to form embryoid bodies (EBs). Day 0 differentiation media is composed of N2B27 [Advanced DMEM F12 (GIBCO, #12634010), Neurobasal (GIBCO, #21103049) vol:vol], supplemented with N2 (Life Technologies, #17502048), B27 without Vitamin A (Life Technologies, #12587010), penicillin/streptomycin 1% (Gibco, #15070-063), β-mercaptoethanol 0.1% (Life Technologies, #31350010), L-Glutamine (200 mM, Life Technologies)] supplemented with ROCK inhibitor Y-27632 (10μM, Stem Cell Technologies, #72304), CHIR99021 (0.7 μM, SELLECKCHEM #S1263), LDN-193189 (200 nM, SELLECKCHEM # S2618), SB-431542 (10 μM, SELLECKCHEM # S1067), SAG (500 nM, SHH agonist, SIGMA, #566660). Differentiation media was changed every other day; EBs were collected and centrifuged at low speed (120 g) to avoid aggregation and replated in the same wells with new media. On Day 2 of differentiation, ROCK inhibitor Y-27632 was withdrawn. On day 4, cells were exposed to a CHIR-BOOST: CHIR-99021 7 μM until day 8 to ensure midbrain-hindbrain barrier identity, then CHIR-99021 3 μM until day 10 when withdrawn [2]. On day 6, LDN, SB, and SAG were withdrawn. On day 10, media was changed to mDA media [N2B27 supplemented with BDNF (brain-derived neurotrophic factor, 10ng/ml; Peprotech #450-02), GDNF (glial cell line-derived neurotrophic factor, 10 ng/ml; Peprotech # 450-10), TGFb3 (transforming growth factor type b3, 1 ng/ml; Life technologies, # PHG9305)] complemented with Notch inhibitor DAPT (10μM, Tocris, #2634) to ensure terminal neuronal differentiation until day 16. At day 16, DAPT was withdrawn, and EBs were cultured in mDA media.

### hiPSC diCerentiation into midbrain Astrocytes (mAstro)

For hiPSC differentiation into midbrain astrocytes, we adapted a protocol developed by others [19], and ensured midbrain/hindbrain identity by using CHIR-BOOST for mDA differentiation [2]. The differentiation protocol was similar to the mDA differentiation protocol described, with different culture media. Day 0 differentiation media comprised N2B27 supplemented with ROCK inhibitor Y-27632 (10 μM, Stem Cell Technologies, #72304) and CHIR99021 (0.7 μM, SELLECKCHEM #S1263). On Day 2 of differentiation, ROCK inhibitor Y-27632 was withdrawn. On day 4, cells were transferred to mAstro media [N2B27 supplemented with CNTF (5 ng/ml; Peprotech #450-13) and FBS (2%, Life Technologies, A31665-01)] and exposed to a CHIR-BOOST: CHIR-99021 7 μM until day 8 to ensure midbrain-hindbrain barrier identity, then CHIR-99021 3 μM until day 10 when withdrawn [2]. Then EBs were cultured in the mAstro media.

### 2D monolayer cultures in 96-well plates

For 2D monolayer cultures, on day 25, EBs were dissociated to single cells using gentle enzymatic dissociation [Papain, 37 °C, 20min (Worthington, #LS003126) or TrypLE RT, 15min (Gibco, #12563-011)] and plated in optically clear flat-bottom 96-well plates (PhenoPlate 96-well, REVVITY HEALTH SCIENCES INC, # 6055302). For monocultures, mDA neurons were plated at a density of 35K/well, or mAstro plated at 5K/well, and cultivated in mDA media and mAstro media, respectively. For mDA-mAstro cocultures, mDA and mAstro were plated together at a density of 30K/Well and 3K/Well, respectively, and cultivated in mDA media and mAstro media, respectively. To ensure even distribution, 96-well plates were centrifuged at 120xg for 2 min. After plating at day 25, the media was changed bi-weekly.

### 3D organoids and 3D mDA-mAstro assemboids

For 3D organoid and assembloid cultures, at day 25, EBs were left undissociated and transferred to Ultra-low attachment Round bottom 96-well plates (Corning, #7007). For individual organoids, mDA EBs or mAstro EBs were plated at a density of one EB/well and cultivated in mDA media and mAstro media, respectively. For mDA-mAstro assembloids, mDA and mAstro EBs were plated together at a density of one EB/well each and cultivated in one-to-one mDA-mAstro media. After plating at day 25, the media was changed bi-weekly.

### Chemically-induced human mDA neurodegeneration models (CBE and rotenone)

For chemically induced neurodegeneration models using CBE (SIGMA, # 234599100MG) or Rotenone (Thermo Fisher Scientific, # AC132370050), 2D monolayer cultures were treated at day 35 with CBE [0.5mM-5mM] or Rot [0.1µM-10µM]. To test for neuroprotection, AGTR1 inhibitors [Candesartan, Thermo fisher scientific, # AAJ62818MC; Telmisartan, Thermo fisher scientific, # AAJ61441MD; Valsartan, Thermo fisher scientific, # AC465560010] were added 4-6h after CBE- or Rot-induced neurodegeneration. The 96-well plates were kept at 37^0^C and 5% C02 and imaged on ImageXpress Confocal HT.ai High-Content Imaging System (Molecular Devices) at the Center for Advanced Light Microscopy (CALM, UCSF) at days 35, 38, and 42.

### Automatic counting using Cell Profiler + FCS Express 7.0 technology

We used Cell Profiler 4.0 (**Error! Hyperlink reference not valid.** [20]) to segment and count the total cell number per well and quantify TH-TdTom+ cell numbers. To ensure impartial quantification of TH-TdTom+ cells, we employed FCS Express 7 software (De Novo Software), allowing us to visualize cells in FACS plots and eliminate doublets and debris from the counting process.

### Immunohistochemistry

For 2D monolayer cultures, cells in 96-well plates were fixed in 4% paraformaldehyde (PFA) (AOymetrix #MFCD00133991) in DPBS for 10 min at room temperature. For immunostaining of 3D organoids and assembloids, organoids/assembloids were in 4% paraformaldehyde (PFA) (AOymetrix #MFCD00133991) in DPBS for 30 min at 4^0^C. Then, the organoids were embedded in OCT and cut into 10 µm slices using a cryostat. Sections were permeabilized with 0.2% Triton X-100 and blocked with 2% Normal Donkey Serum (Jackson Labs, #AB_2337258) in DPBS. The samples were subsequently incubated with the primary antibody overnight at 4 °C. The next day, after washing with DPBS, the samples were incubated with secondary antibody conjugated with Alexa Fluor 488- 555-, or 647- (Thermo Fisher) diluted at 1:2000 in DPBS + 0.2% Triton X-100 + 2% Normal Donkey Serum for 1 hour at room temperature. Then the samples were washed with DPBS and counterstained with 40, 6-diamidino-2-phenylindole (DAPI) (Sigma, #D9542). Images were visualized using an ImageXpress Confocal HT.ai High-Content Imaging System (Molecular Devices). Rabbit anti-TH (1:800, PelFreez and 1:1000, Immunostar), goat anti-FOXA2 (1:500, R&D), Rabbit anti-AGTR1 (1:2000, Alomone Labs), Mouse anti-GFAP (1:1000, Sigma), mouse anti-alpha-Syn (1:800,BD Bioscience) and Rabbit Phospho-α-Synuclein (Ser129) (1:1000, D1R1R, Cell signaling) were used for immuno-fluorescent staining. Donkey anti-mouse, goat, rabbit, or chicken secondary antibodies conjugated with Alexa Fluor-488, Alexa Fluor-555, or Alexa Fluor-647 fluorophore (1:2000, Life Technologies) were used. Nuclei were counterstained with DAPI (Sigma, #D9542).

### RNA extraction and Real-time qRT-PCR

Total RNAs from samples were isolated with the RNeasy Micro Kit (QIAGEN, #74004). 1 μg of RNA was used to generate cDNA using the iScript Reverse Transcription Supermix (BioRad, #170 8841). Real-time qRT-PCR was performed using the High-Capacity cDNA Reverse Transcription kit (Thermo Fisher Scientific, #4368814) in a Bio-Rad CFX96 Thermal Cycler. qRT-PCR reactions were performed using the Taqman Fast Advanced Master Mix (Thermo Fisher Scientific, #4444556) on a QuantStudio™ 7 Flex Real-Time PCR System, 384-well (Thermofisher). All reactions were performed according to the manufacturer’s protocol. Primers were ordered from predesigned Taqman PCR assays (Thermofisher) with the following catalog numbers: (GAPDH: Hs02786624_g1; SNCA: Hs00240906_m1; SNCB: Hs01015410_g1; SNCAIP: Hs00917422_m1). Results were normalized to GAPDH.

### Bulk RNA-sequencing

RNA-seq library preparation, sequencing, alignment, and gene expression matrix were performed and delivered by BIOSTATE.ai, a cutting-edge RNA sequencing technology. Differential gene expression was performed using DESeq2 (v. 1.12.4) [21].

### Analysis of RAAS pathway gene expression using scRNAseq and snRNAseq databases

Available sc/snRNA-seq data were analyzed from recent human and mouse high-resolution snRNA-seq datasets [22,23] using the Allen Brain Cell Atlas website (https://knowledge.brain-map.org/abcatlas). Human aged midbrain snRNAseq [5] data were analyzed using the https://singlecell.broadinstitute.org/ website with reference provided in the study. Human Protein Atlas (https://www.proteinatlas.org/) scRNA-seq data [24] were also used for analysis.

### CRISPRi knockdown of AGTR1

The iFluo-gRNA-dCas9-KRAB-EGFP vector, designed for the co-expression of dCas9-KRAB, EGFP, and dual gRNAs, was constructed using components from the Lenti-dCas9-KRAB-blast vector (Addgene #89567) and the CROP-seq-opti vector (Addgene #106280). EGFP was inserted in place of the Blasticidin S deaminase (BSD) gene during vector assembly. To achieve CRISPRi-mediated knockdown of AGTR1, two pairs of gRNAs targeting regions within ±100 bp of the transcription start site (hg38: chr3:148697834(-), chr3:148697935(+); and chr3:148697848(-), chr3:148697984(+)) were designed using GuideScan [25] and CHOPCHOP [26]. These gRNAs were predicted to have no oO-target sites with zero or one mismatch, 0–2 sites with two, and 0–7 sites with three mismatches. The dual gRNAs were cloned into the iFluo-gRNA-dCas9-KRAB-EGFP vector as previously described [27].

For lentiviral packaging, 7.5 μg of the gRNA plasmid was co-transfected with 1.5 μg of pMD2.G (Addgene #12259) and 4.5 μg of psPAX2 (Addgene #12260) into 293 T-LentiX cells (Takara Bio #632180) seeded in a T-75 flask, using PolyJet transfection reagent (SignaGen #SL100688). The transfection reagent was removed 8–14 hours post-transfection, and viral supernatants were collected every 24 hours for three consecutive days. The harvested virus was filtered through a 0.45 μm membrane and concentrated using Amicon Ultra-15 Centrifugal Filters (Millipore #UFC901024). Viral titers were estimated by measuring the percentage of GFP-positive cells among the total cell population following infection.

During the preparation of this work, the author(s) used Grammarly and Chat-GPT in order to improve the clarity and readability of the text. After using this tool/service, the author(s) reviewed and edited the content as needed and take(s) full responsibility for the content of the publication.

## RESULTS

### Development of a Scalable High-Content Screening Platform for hiPSC-Derived mDA Neurons

To elucidate the neuroprotective mechanisms and identify disease-modifying therapeutics for mDA neurons in human models of PD, we established a high-content screening platform utilizing hiPSC-derived mDA neurons. We employed the KOLF2.1J-TH-TdTom hiPSC line [15,16] for live imaging of mDA neurons. To ensure high purity and reproducibility, we refined a midbrain DA differentiation protocol incorporating a CHIR-boost technique to specify mDA neurons corresponding to the SNpc [2]. We adapted and improved this protocol, originally in a 2D monolayer, to produce 3D embryoid bodies (EBs), allowing a more reproducible and improved ratio of TH-neurons. After terminal differentiation (D25), mDA neurons were dissociated and plated at low density in 96-well plates (**Fig. 1A-C**). Our optimized protocol consistently produced ∼80% TH-TdTom+ mDA neurons in a 96-well format (**Fig. 1C-D**). A strong correlation between TH-TdTom fluorescence and TH-antibody staining was observed, validating the TH-TdTom CRISPR knock-in model (**Fig. 1E**). Immunofluorescent labeling further verified co-expression of critical mDA markers, including FOXA2 (midbrain floor plate and mDA marker), TH protein, and AGTR1 (**Fig. 1E-F**). 3D EBs were kept in their 3D shape to establish a 3D mDA organoid model, where we confirmed expression of TH-TdTom throughout the organoid, correlated with TH-Antibody and AGTR1 (**Fig. 1G**).

**FIGURE 1.**
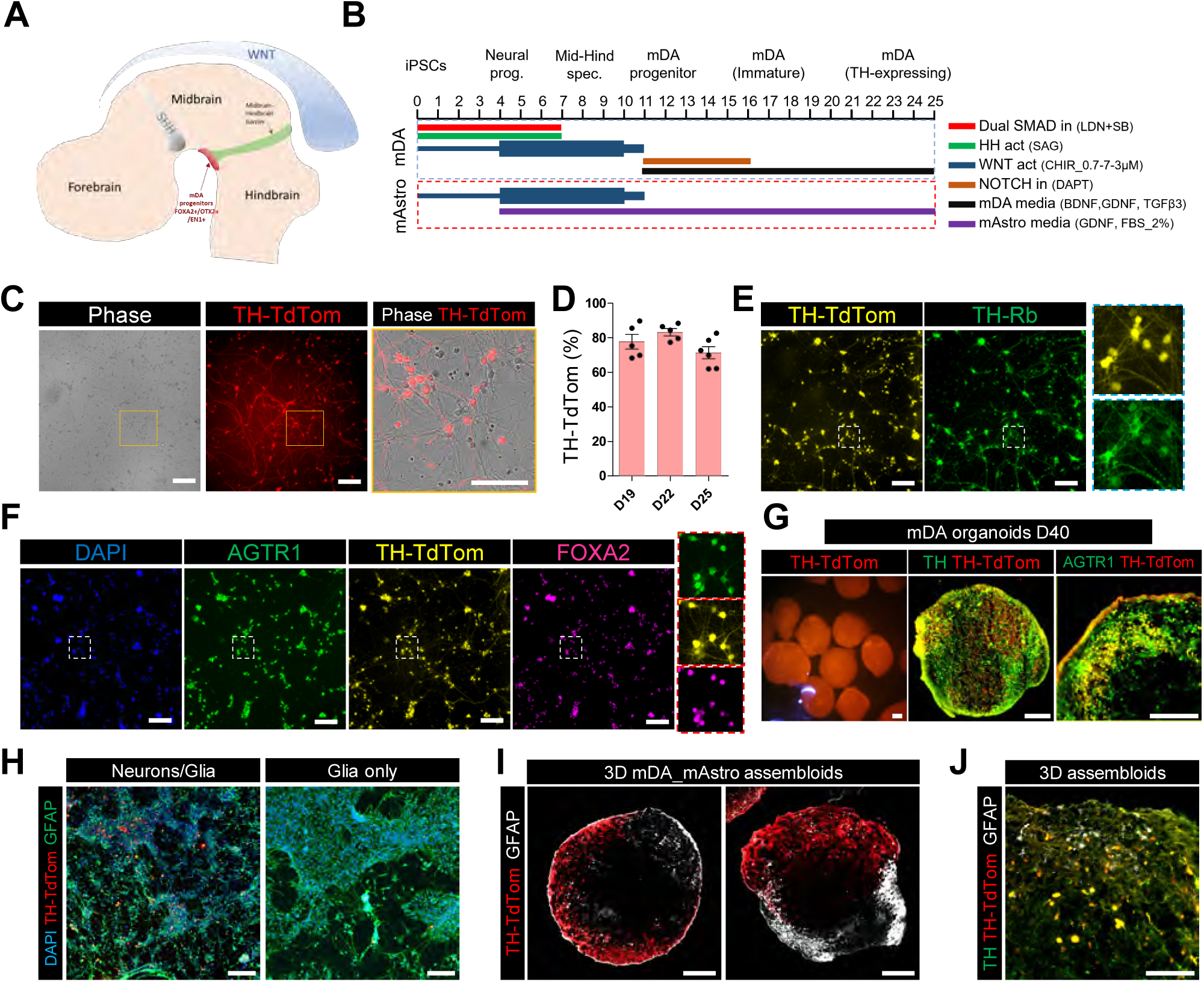
DiCerentiation of hiPSCs into midbrain dopamine neurons and astrocytes in 2D and 3D cultures. (**A**) A schematic diagram shows midbrain dopamine (mDA) progenitors near the midbrain-hindbrain boundary. (**B**) A summary of the protocols used to differentiate hiPSCs to mDA neurons and midbrain astrocytes (mAstro). mDA neuron organoids or mDA-mAstro neuron-glia assembloids were formed by assembling embryoid bodies at D25. (**C, D**) The TH-TdTom hiPSC line, generated via CRISPR knock-in, was used to visualize mDA neurons. mDA cultures are ∼80% TH-TdTom+ (n=6). (**E**) TH-TdTom reporter expression correlates with TH antibody staining. (**F**) mDA neurons co-express mDA progenitor marker FOXA2 and AGTR1. (**G**) D40 3D mDA organoids show robust TH-TdTom, TH, and AGTR1 expression. (**H**) Images of the neuron-glia 2D co-culture system. (**I**) D60 mDA-mAstro assembloids display distinct TH-TdTom+ and GFAP+ regions. (**J**) GFAP+ glia successfully infiltrate the TH-TdTom+ neuronal region. Scale bars: 100 μm (C-F,H,J); 200 μm (G,I).

To determine whether proper midbrain patterning was achieved in our 2D and 3D cultures, we evaluated region-/cell-type-enriched markers as previously used [2]. We confirmed the expression of TH, FOXA2, LMX1A, EN1, OTX2, AXIN2, FOXA1, SHH, and MSX1, with low to absent expression of PAX6, NKX2.2/2.1, POU5F1, HOXA2, HOXB1, and DBX1 (**Fig. S1A-F**). Quantifications on mDA organoids at D40 showed that 85% of TH-TdTom+ population expressed AGTR1, while 65-80% of AGTR1+ population expressed TH, depending on the TH-TdTom hiSPC clones used (**Fig. S1G-H**). Moreover, we found a high correlation between TH-TdTom and AGTR1 expression (r^2^=0.588; **Fig. S1I**).

The prepropeptide ligand in the RAAS pathway, angiotensinogen (AGT), is highly expressed in astrocytes [28]. To enrich our culture model and investigate neuron-glia interactions, we developed a 2D midbrain neuron-astrocyte co-culture system incorporating hiPSC-derived midbrain astrocytes (mAstro), patterned using a CHIR-boost protocol we adapted from previous published reports [2,19] to confer midbrain/hindbrain identity (**Fig. 1B**). In these cultures, TH-TdTom+ mDA neurons were surrounded by GFAP+ astrocytes (**Fig. 1H**). Additionally, we assembled mDA and mAstro EBs individually to create a 3D midbrain neuron-glia (mDA-mAstro) assembloid model, exhibiting a TH-TdTom+ mDA side opposite to a GFAP+ mAstro side (**Fig. 1I**). Moreover, after 60 days of maturation, GFAP+ glia successfully infiltrated the TH-mDA side, showing the robustness of our coculture model (**Fig. 1J, Fig. S1J**).

Together, our hiPSC-derived mDA neuron platform achieves high purity of SNpc-like TH-TdTom+ mDA neurons in 2D mDA, 2D mDA-mAstro neuron-glia co-culture, 3D mDA organoid, and 3D mDA-mAstro neuron-glia assembloid models, enabling subsequent investigations of mDA neuron differentiation and survival.

### Analyses of high-resolution snRNA-seq data identify the expression of RAAS pathway genes in the human nigrostriatal pathway, distinct from that of the mouse

To investigate the role of RAAS signaling in human PD, we first examined the expression of RAAS components in human and mouse brains using databases composed of high-resolution single-cell/single-nuclei RNA sequencing (scRNA-seq/ snRNA-seq) and spatial antibody-based profiling data [5,22–24]. The main components of the RAAS pathway (**Fig. 2A**) include the prepropeptide ligand AGT, which is processed by two proteases, Renin (REN) and angiotensin converting enzyme (ACE), into the active ligand Angiotensin II (Ang II). Ang II activates AGTR1, a Gq-coupled GPCR that activates a phosphatidylinositol-calcium second messenger system. Ang II can also activate AGTR2, which couples to Gi, leading to the inactivation of the extracellular signal-regulated kinases (ERK) pathway and often opposing the effects of AGTR1 [29–31].

**FIGURE 2.**
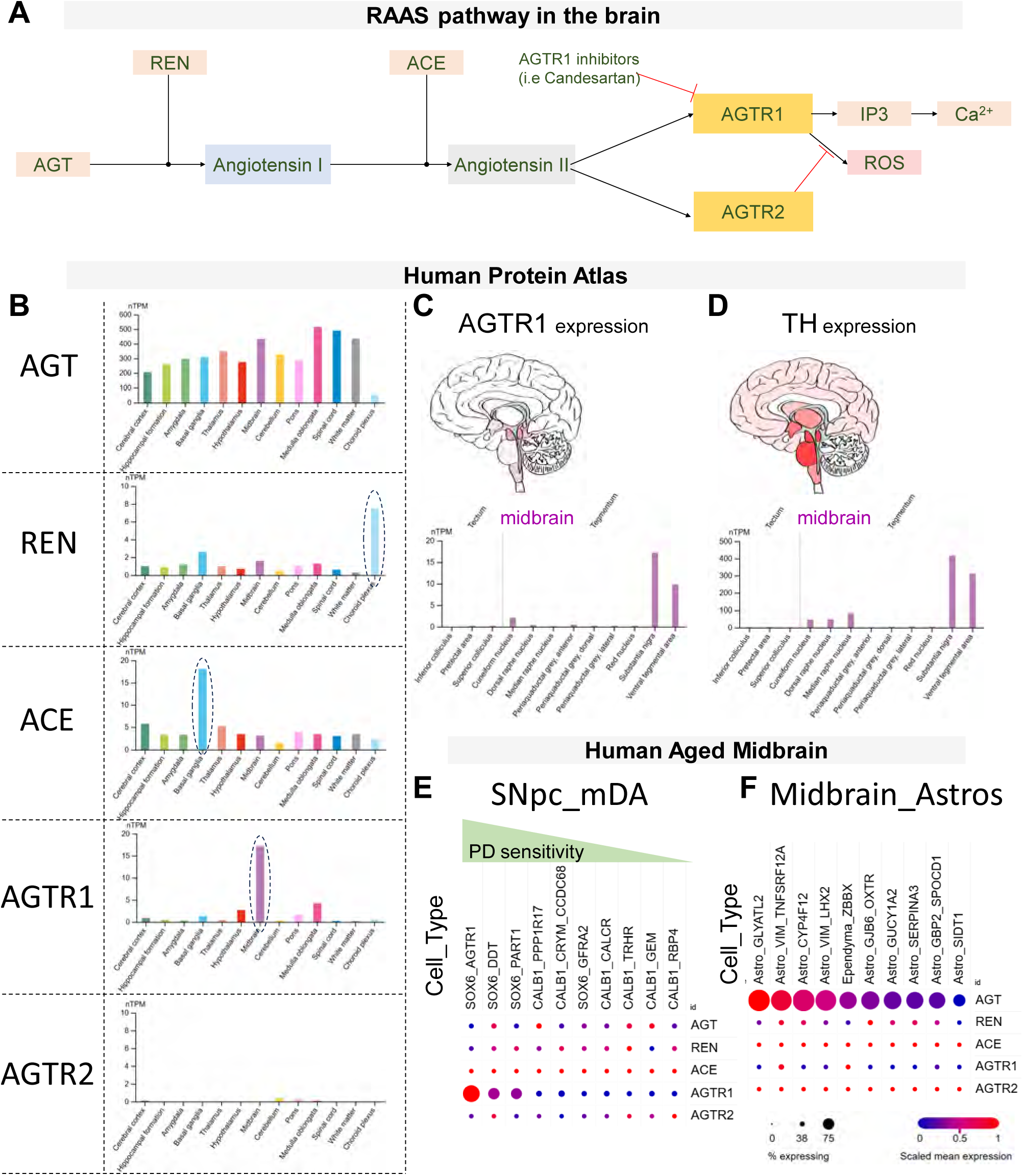
RAAS Pathway Component Expression in the Human Brain. (**A**) A diagram of the major RAAS pathway components in the human brain. AGT, angiotensinogen; REN, renin. (**B-D**) Graphs from the Human Protein Atlas show broad expression of AGT, enriched REN in choroid plexus cells with detectable levels in the basal ganglia, ACE enrichment in the basal ganglia, AGTR1 enrichment in the midbrain, particularly in the substantia nigra and ventral tegmental area, mirroring TH expression (**C-D**), and little to no AGTR2 expression. (**E, F**) Data from the Human Aged Midbrain snRNAseq database show low AGT, REN, ACE, and AGTR2 in SNpc mDA neurons, high AGT in midbrain astrocytes, and AGTR1 expression in the SNpc populations most vulnerable in PD.

Using the Human Protein Atlas, we found that AGT was widely expressed across brain regions, consistent with its expression in the astrocytes that are widely distributed in the brain. REN expression was primarily limited to choroid plexus cells, with lower levels in the basal ganglia, while ACE was predominantly expressed in the basal ganglia (**Fig. 2B**). Notably, AGTR1 expression was highly enriched in a subset of TH-positive neurons in the midbrain, particularly in the SNpc (**Fig. 2B-D**). The SnRNA-seq data from aged human midbrains further confirmed AGTR1 expression in a subset of PD-vulnerable SOX6+ SNpc DA neurons [5], while AGT was expressed in the midbrain astrocytes. REN, ACE, and AGTR2 showed minimal or no expression in SNpc mDA neurons or midbrain astrocytes (**Fig. 2E,F**).

The SNpc DA neurons connect to medium spiny neurons (MSNs) in the striatum [32]. Using whole-brain human and mouse snRNA-seq data [22,23], we identified SNpc DA neurons by their expression of TH, DAT/SLC6A3, FOXA2, and EN1 [2], and MSNs by DARPP-32, DRD1, and DRD2 [33] (**Fig. S2A-B; Fig. S3A-B**). In the human brain, we confirmed AGT expression primarily in astrocytes and identified REN expression in choroid plexus cells and MSNs. ACE expression was also prominent in MSNs. AGTR1 expression was limited to specific neuronal subsets, including PD-vulnerable TH⁺SOX6⁺CALB1⁻ SNpc neurons [5], with minimal expression in MSNs. In agreement with the Human Protein Atlas data, no to little expression of AGTR2 was detected in the human brain (**Fig. S2C-D**).

Comparative snRNA-seq analysis of human and mouse brains revealed species-specific differences. In humans, the PD-vulnerable TH⁺SOX6⁺CALB1⁻ DA neuron subtype expressed AGTR1 but not AGTR2, whereas the corresponding mouse population lacked AGTR1a/b and expressed AGTR2 (**Fig. S2C-D; Fig. S3C-D**). Given AGTR1’s pro-degenerative role and AGTR2’s neuroprotective role in DA neurons [34], these differences highlight the limitations in using mouse models to study AGTR1-mediated mechanisms, emphasizing the value of human-based *in vitro* platforms.

Together, these findings establish the expression of RAAS components in the human nigrostriatal pathway, with AGTR1 expression in PD-vulnerable SNpc DA neuronal subtypes and AGT expression by surrounding astrocytes. Moreover, we identify choroid plexus cells and MSNs as key sources of REN, the rate-limiting protease in producing the active ligand Ang II.

### Chemically Induced PD-Relevant Neurodegeneration Models

To model PD *in vitro* using our scalable high-content platform with hiPSC-derived mDA neurons, we developed two chemically induced neurodegeneration models in which we observed significant loss of the TH-positive mDA neurons (**Fig. 3**). We chose chemical approaches rather than genetic means (e.g., patient-derived or genetically edited iPSC lines) because genetic methods often result in little or no neurodegeneration in culture settings without stress challenges [35–40].

**FIGURE 3.**
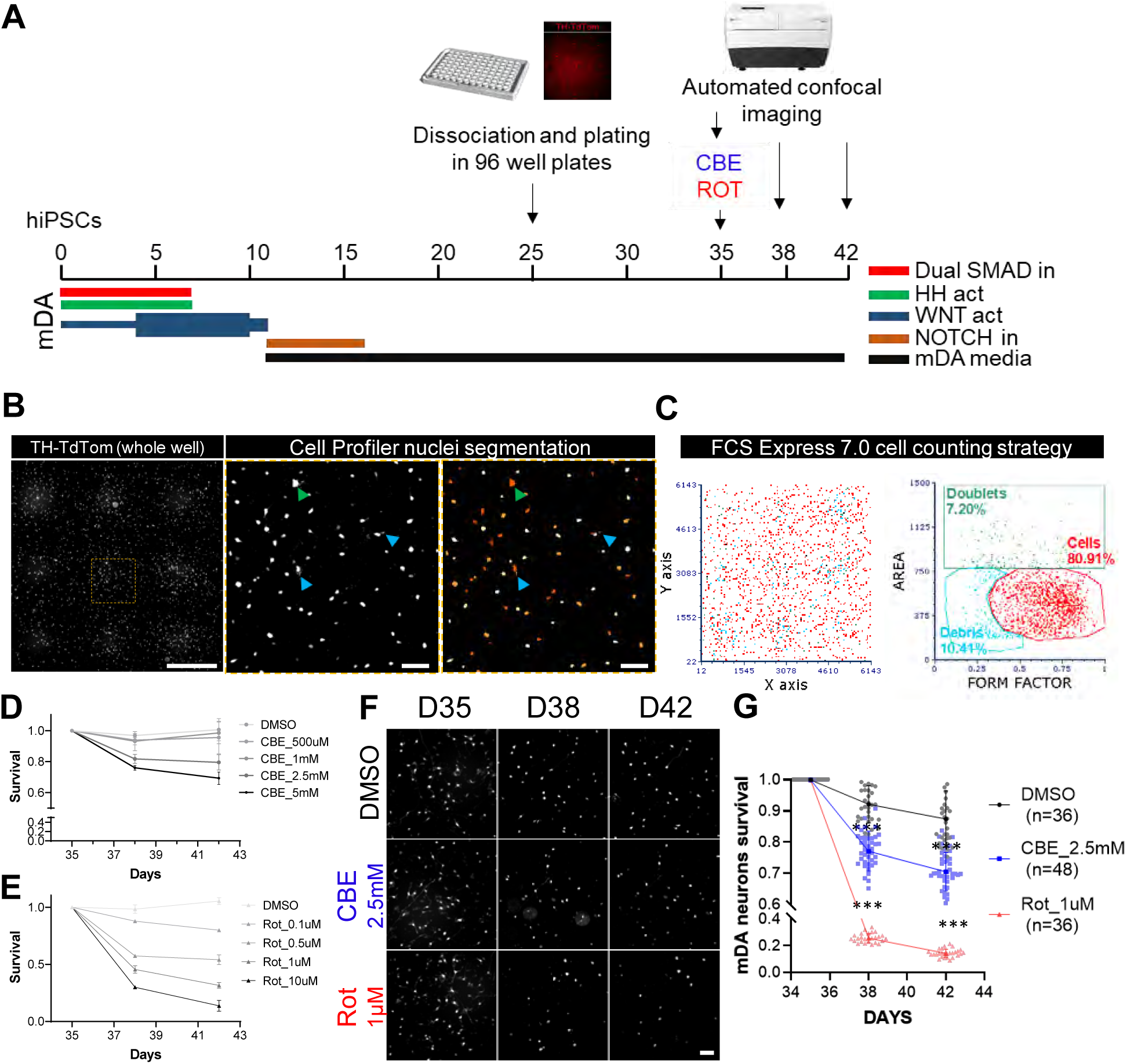
Chemically induced neurodegeneration models using CBE and Rotenone in hiPSC-derived mDA 2D cultures. (**A**) Diagram of the survival experimental pipeline. mDA neurons are treated with CBE vs Rotenone (Rot) at D35 and imaged by automated confocal microscopy at D35, D38, and D42. (**B, C**) mDA segmentation using Cell Profiler 4.0; FCS Express 7.0 image cytometry allows the removal of doublets (green dots and arrows) and debris (blue dots and arrows) from the counting. (**D, E**) To induce neurodegeneration, CBE [500 μM-5 mM] and Rot [0.1 μM -10 μM] were used (n=5 for each condition). (**F**) [CBE] at 2.5 mM and [Rot] at 1 μM were further selected to model a moderate (20-30%) and more pronounced (50-70%) loss of mDA neurons. (**G**) [CBE]_2.5 mM or [Rot]_1 μM treatment on a large sample size shows high reproducibility. *** p<0.001. Scale bars: 1mm (B, left); 100 μm (B, right, F)

A key challenge in hiPSC-derived neuronal cultures is neuronal clumping, which hinders accurate cell counting. We optimized cell density, substrate coating, and dissociation timing (**Fig. S4A-D**). Delaying dissociation to day 25 (D25) from day 11 (D11) significantly reduced clumping (**Fig. S4A-B**). Additionally, we developed a custom polyethylenimine (PEI) plus laminin (Lam) coating, which outperformed the standard poly-D-lysine (PDL) plus laminin coating, maintaining isolated neurons up to D60 and improving the accuracy of automated counting (**Fig. S4C-D**). This optimization enabled long-term, clump-free cultures with stable TH-TdTom expression by D35, the stage selected for initiating drug treatment and survival studies via live-imaging (**Fig. 3A**). Whole-well imaging, coupled with automated nuclei segmentation based on TH-TdTom fluorescence and analysis by image cytometry, allowed precise fluorescence intensity analysis, effectively excluding doublets and debris for accurate mDA neuron quantification (**Fig. 3B,C**). Representative nuclei segmentation and automated counting over 14 days were shown in **Fig. S4E-F**, with the same strategy applied to 3D assembloid slice counting (**Fig. S4G**).

We treated hiPSC-derived mDA neurons with conduritol B-epoxide (CBE), an inhibitor of glucocerebrosidase (GBA1) associated with PD risk [17], and Rotenone (Rot), a mitochondrial complex I inhibitor that induces neuronal death [18]. CBE treatment triggered dose-dependent mDA neuron loss starting at 2.5 mM (**Fig. 3D,F**), while Rot induced neuronal loss from 0.1 µM (**Fig. 3E-F**). Time-course imaging of neuronal loss with increasing CBE and Rot concentrations was shown in **Fig. S5A-B**. We selected 2.5 mM CBE and 1 µM rotenone to achieve moderate (20–30%) and profound (50–70%) mDA neuron loss, respectively (**Fig. 3F-G**). The reproducibility of these models was validated across large sample sizes and two independent clones of the KOLF2.1J-TH-TdTom cell line, demonstrating consistent results (**Fig. 3G; Fig. S5C-D**).

These chemically induced neurodegeneration models, the first to combine PD-relevant neurotoxic agents with hiPSC-derived mDA neurons in a 96-well format, enable quantitative and scalable testing of neuroprotective compounds, offering a robust platform to identify therapeutic candidates for PD.

### Neuroprotective ECects of AGTR1 Inhibition

Next, we evaluated the effects of AGTR1 inhibition on both 2D and 3D hiPSC-derived mDA neurodegeneration models (**Fig. 4A-B**). Since astrocytes play critical roles in supporting neuronal functions [41], we used 2D and 3D mDA-mAstro co-cultures and ensured accurate cell counting by leveraging TH-TdTom fluorescence specific to mDA neurons.

**FIGURE 4.**
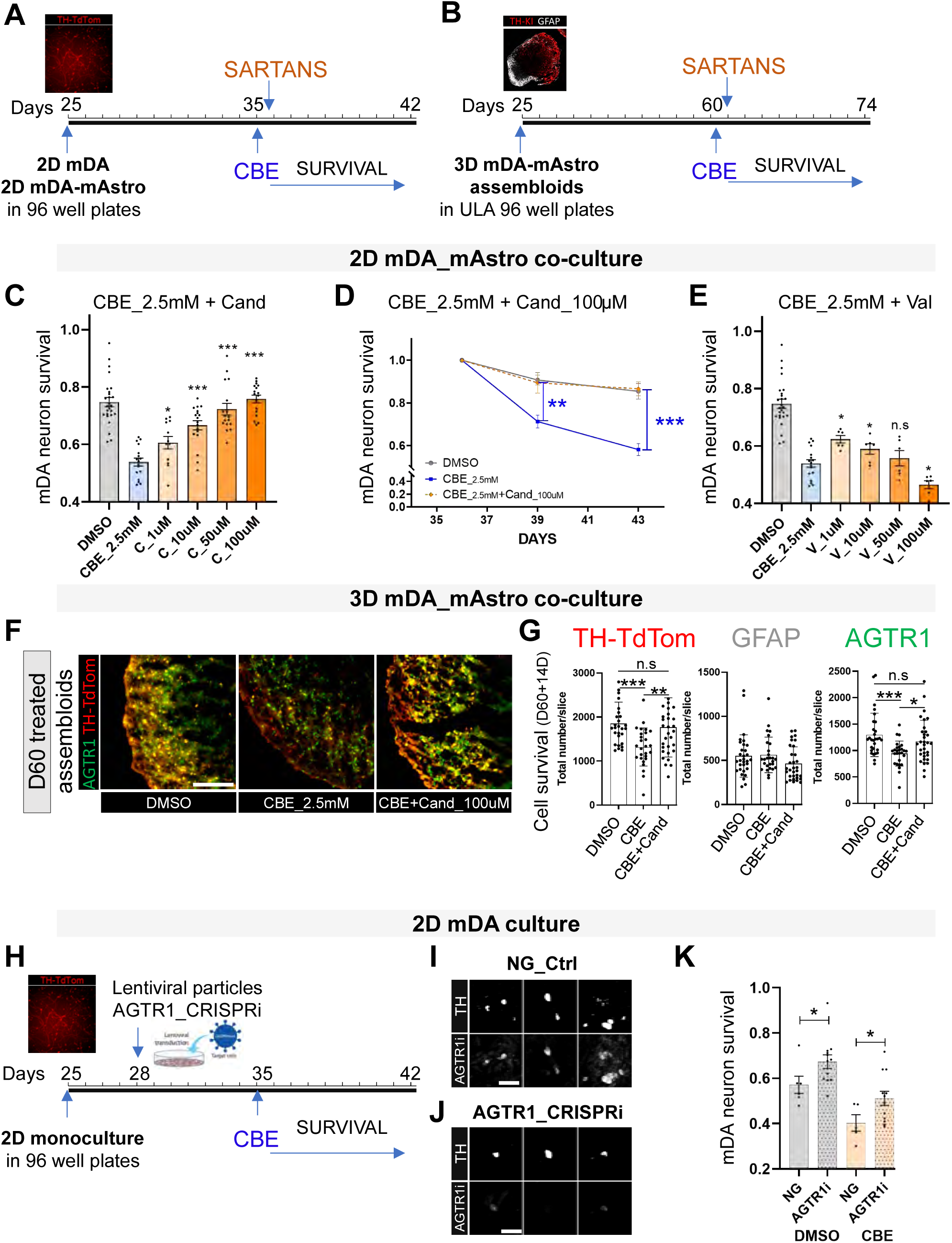
AGTR1 inhibition confers neuroprotection in CBE-induced 2D and 3D human mDA neurodegeneration models: (**A, B**) Pipeline of survival experiments in 2D monolayer and 3D mDA-mAstro assembloids; CBE treatment was followed with AGTR1 inhibitors (sartans). (**C**) Increasing concentrations of Candesartan (1μM-100μM) were added to the mDA neurons a few hours after CBE_2.5mM treatment, showing a dose-dependent protection against neuronal death (D42 compared to D35). (**D**) Using Cand_100μM, we found a nearly total neuroprotection against CBE at D38 and D42. (**E**) Valsartan showed neuroprotection at low doses (1-10μM), albeit neurotoxic at higher doses. (**F, G**) In 3D mDA-mAstro assembloids treated at D60 for 14 days, Candesartan protects TH-TdTom- and AGTR1-expressing populations against CBE-induced cell death while GFAP+ glia remain unchanged. (**H**) Strategy for lentiviral delivery of AGTR1-CRISPRi machinery in 2D mDA cultures. (**I,J**) AGTR1-CRISPRi knockdown was confirmed by immunostaining 1-week post-infection. (**K**) Protection of mDA neurons in DMSO and CBE conditions was observed in the AGTR1i conditions. * p<0.05, **p<0.01, *** p<0.001. Scale bars: 100 μm (F); 30 μm (I, J)

We first tested the AGTR1 inhibitors candesartan and valsartan post-CBE treatment in 2D mDA and astrocyte cocultures. Sartans are known to be highly protein-bound [42], significantly reducing free drugs available for binding to AGTR1. Therefore, we tested concentrations of these sartans, ranging from 1 μM to 100 μM. Candesartan exhibited dose-dependent neuroprotection, achieving near-complete protection against CBE-induced mDA neuron loss at 100 µM by days 38 and 42 (**Fig. 4C-D**). Valsartan also provided neuroprotection at low doses but displayed toxicity at higher concentrations (**Fig. 4E**). We next tested the effect of candesartan in the 3D mDA-mAstro assembloid model (**Fig. 4B**). Candesartan preserved TH+ and AGTR1+ neurons against CBE-induced loss after 14 days without affecting GFAP+ astrocytes (**Fig. 4F-G**).

To genetically inhibit AGTR1, we developed an AGTR1-CRISPRi system delivered via lentiviral vectors, validated by immunostaining 7 days post-infection (**Fig. 4H-J; Fig. S6**). AGTR1-CRISPRi conferred significant neuroprotection in both DMSO and CBE-treated conditions, though partial transduction limited knockdown efficiency, resulting in incomplete protection compared to candesartan (**Fig. 4K, Fig. S6B-E**).

Against rotenone (Rot)-induced neurotoxicity, AGTR1 inhibitors showed limited efficacy at the high Rot dose (1 µM) due to severe neurodegeneration. However, at the lower Rot dose (0.2 µM), valsartan provided partial neuroprotection at high concentrations (50–100 µM) in 2D cultures, while candesartan did not show neuroprotection, contrary to CBE-treated conditions (**Fig. S7A-C**). In mature 3D mDA-mAstro assembloids (D80; **Fig. S7D**), even low Rot doses (0.2 µM) exhibited near-total TH-TdTom loss (**Fig. S7E**); high Rot toxicity likely accounted for undetectable effects of candesartan and valsartan (**Fig. S7F**).

Together, these results demonstrate significant neuroprotection by AGTR1 inhibition, particularly Candesartan, in CBE-induced 2D and 3D models, with more limited efficacy against Rot-induced damage due to its high toxicity leading to severe neurodegeneration.

### Transcriptomic Analyses of mDA neurons upon AGTR1 Inhibition

To elucidate the mechanisms underlying AGTR1 inhibition-mediated neuroprotection, we performed bulk RNA sequencing in hiPSC-derived mDA 2D model treated with or without CBE, and with or without AGTR1 inhibition. RNAs were collected 18 hours post-treatment before overt cell death, to minimize cell death-related gene profiles (**Fig. 5A**). We focused on three AGTR1 inhibitors: candesartan, valsartan, and telmisartan (thereafter sartans) to explore neuroprotective pathways mediated by AGTR1 inhibition.

**FIGURE 5.**
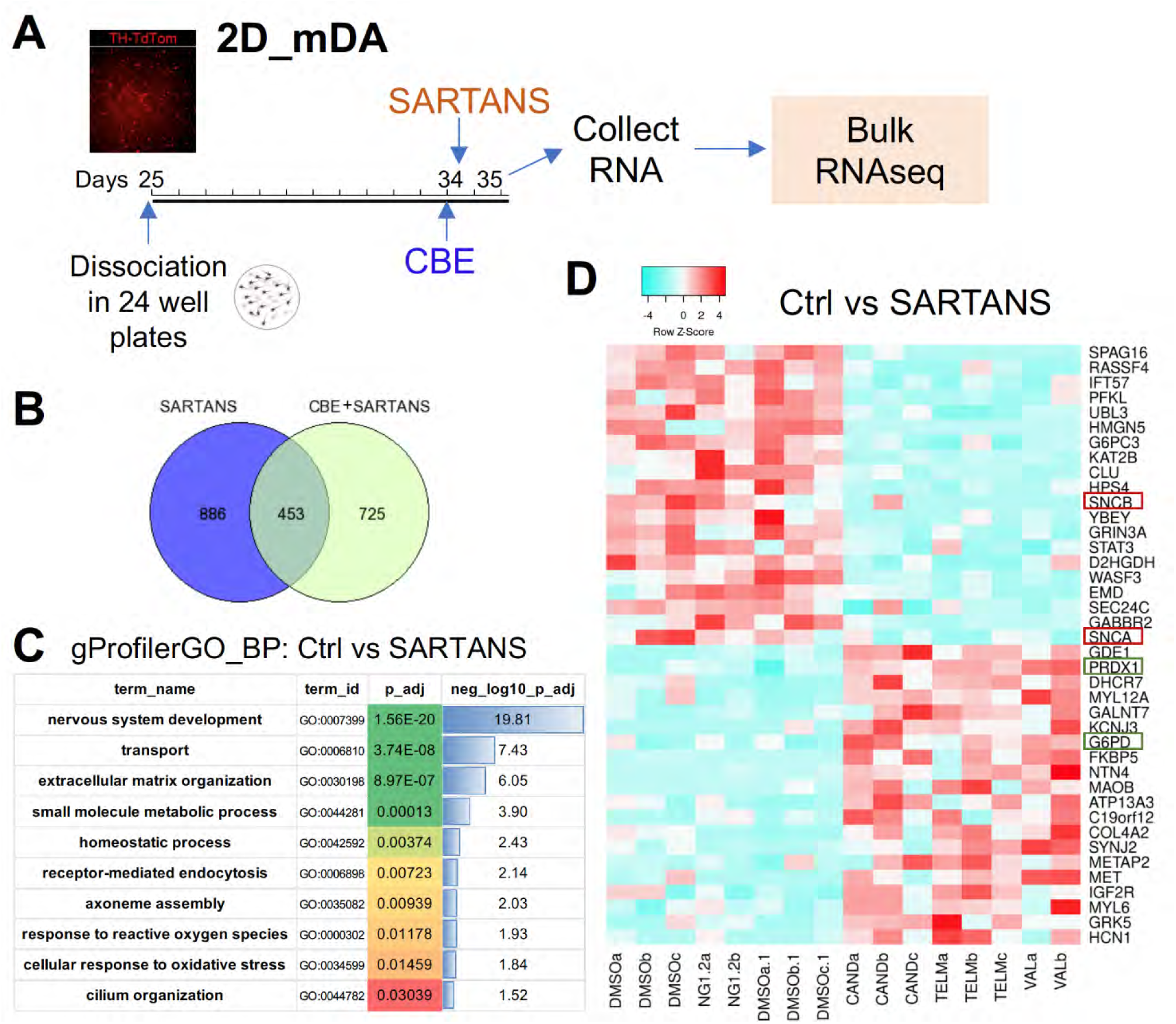
Transcriptomic analysis to discover gene expression changes associated with AGTR1 inhibition. (**A**) The schematic shows the culture pipeline for RNA collection and bulk RNA-seq. (**B**) Venn diagram showing the common differentially expressed genes in Sartans and CBE+Sartans conditions. (**C**) GO enriched terms in DMSO vs Sartans. Biological processes like transport, endocytosis, and response to ROS were significantly altered by AGTR1 inhibition. (**D**) Top 20 UP and DOWN-regulated genes in DMSO vs Sartans, showing *SNCA* and *SNCB* (red rectangles) as part of the top DOWN-regulated genes and *PRDX1* and *G6PD* (green rectangles) as part of the top UP-regulated genes by AGTR1 inhibition.

Of the 1,339 differentially expressed genes (DEGs) in Ctrl vs Sartan-treated conditions, 453 were consistently altered in CBE + Sartans conditions, forming a candidate gene set for SARTAN-mediated neuroprotection against normal mDA neuron loss in culture and CBE-induced mDA neuronal loss (**Fig. 5B**). Gene Ontology analysis revealed significant enrichment in biological processes, including axonal transport, endocytosis, and response to reactive oxygen species (ROS) (**Fig. 5C**).

Among the top upregulated genes upon AGTR1 inhibition, Peroxiredoxin-1 (*PRDX1)* and Glucose-6-phosphate dehydrogenase *(G6PD)* stood out (**Fig. 5D**), both known to mitigate ROS-induced damage [43–45], suggesting that AGTR1 inhibition may enhance neuroprotection by reducing oxidative stress, as previously proposed [34].

Notably, *SNCA* and *SNCB* were among the top downregulated genes upon AGTR1 inhibition, with *SNCAIP* also downregulated (**Fig. 5D**). *SNCA* encodes α-synuclein, which forms toxic oligomers in Lewy bodies, a hallmark of PD [46], while *SNCAIP* (synphilin-1) promotes α-synuclein aggregation [47]. *SNCB* encodes β-synuclein, which reportedly inhibits α-synuclein aggregation and Lewy body formation [48], though other studies suggest β-synuclein may promote neurotoxicity through α-synuclein-independent pathways [49]. The global downregulation of synuclein-related genes (*SNCA*, *SNCB*, *SNCAIP*) by inhibition of AGTR1 suggests that AGTR1 promotes synuclein expression, directly or indirectly. Inhibiting AGTR1 potentially contributes to neuroprotection by reducing aggregation-prone α-synuclein protein expression.

### AGTR1 inhibition limits toxic phospho-α-Synuclein accumulation in mDA-mAstro assembloid model

To further investigate the link between AGTR1 inhibition and synuclein regulation, we first validated the transcriptional effects of AGTR1 inhibition or activation on synuclein genes via qRT-PCR (**Fig. 6A**). The results confirmed that candesartan treatment significantly reduced *SNCA*, *SNCAIP*, and *SNCB* expression in hiPSC-derived mDA neurons. Conversely, angiotensin II (AngII), an AGTR1 agonist, increased *SNCA* and *SNCB* expression without significantly altering *SNCAIP* (**Fig. 6B-D**).

**FIGURE 6.**
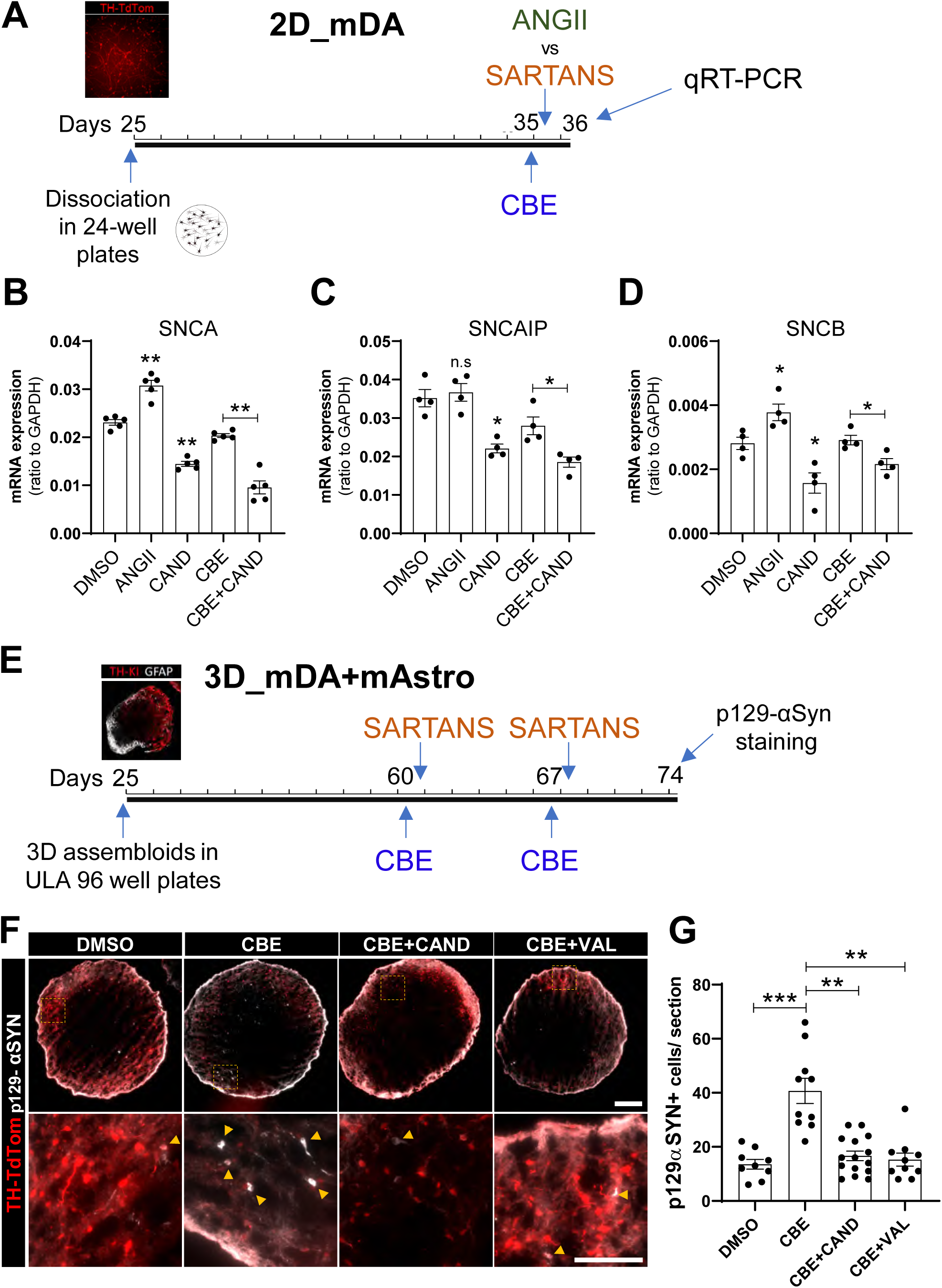
RAAS inhibitors protect against CBE-induced p129 α-synuclein oligomerization. (**A**) The schematic shows different conditions used for qRT-PCR analysis of synuclein gene expression. (**B-D**) qRT-PCR for SNCA, SNCAIP, and SNCB expression using Ang II (AGTR1 agonist) vs Cand (AGTR1 inhibitor). Significance is shown compared to DMSO and CBE + CAND compared to CBE * p<0.05, **p<0.01. (**E**) Pipeline of survival experiments in 3D mDA-mAstro assembloids. CBE treatment followed by AGTR1 inhibitors (sartans) was performed at D60 for 14 days. (**F, G**) CBE treatment increases the number of mDA neurons that are positive for p129-αSYN, a hallmark of PD. Treatment with Candesartan or Valsartan mitigates the number of p129αSYN positive cells. (mDA-mAstro assembloids from 3 independent differentiations). Significance is shown compared to the CBE condition. **p<0.01, *** p<0.001. Scale bars: 200μm (100μm in zoomed images)

To investigate AGTR1 inhibition’s impact on α-synuclein protein levels, we stained 3D mDA-mAstro assembloids with α-synuclein antibody. Despite a strong correlation between TH-TdTom and α-synuclein expression, no apparent differences in α-synuclein protein levels were observed across DMSO, CBE, and CBE + Candesartan conditions (**Fig. S8A**), likely due to the high baseline expression of synucleins in neuronal populations. Indeed, single-cell RNA sequencing data from human whole brain [22] and human aged midbrain [5] confirmed high *SNCA*, *SNCAIP*, and *SNCB* expression in multiple neuronal types, SNpc neurons, with *SNCA* and *SNCAIP* (but not *SNCB*) also highly expressed in medium spiny neurons (MSNs) and SNpc astrocytes (**Fig. S8B-C**).

To specifically assess PD-related pathology, we quantified phosphorylated α-synuclein at serine 129 (p129α-Syn), a marker of pathological aggregates [50], in our 3D mDA-mAstro assembloid model. CBE treatment for 14 days increased p129α-Syn+ TH+ cells, consistent with GBA1-associated PD pathology [51,52] (**Fig. 6E-G**). Notably, candesartan and valsartan significantly reduced p129α-Syn accumulation in TH+ neurons (**Fig. 6F-G**). The reproducibility of CBE-induced p129α-Syn aggregation and its reduction by sartans was validated across multiple mDA-mAstro assembloids from independent differentiations (**Fig. S9**).

Together, these results reveal a novel link between AGTR1 signaling and α-synuclein pathology, in the form of p129α-Syn.

## DISCUSSION

This study established robust "human PD in a dish" models, providing a quantitative, high-throughput approach to induce PD-relevant neurodegeneration in cultured hiPSC-derived midbrain DA neurons. This model enabled us to demonstrate the neuroprotective effects of AGTR1 inhibition on hiPSC-derived mDA neurons. Furthermore, we discovered that AGTR1 inhibition reduces the expression of synucleins, decreasing the accumulation of toxic a-synuclein, underscoring the therapeutic potential of inhibiting AGTR1 for neuroprotection in PD.

### A scalable human cellular platform for modeling PD and identifying therapeutic agents

Our study introduces a robust and highly scalable hiPSC-derived mDA neuron platform that recapitulates PD-relevant neurodegeneration *in vitro*, enabling real-time tracking of mDA neuron survival and high-content screening of neuroprotective molecules. This “PD in a dish” model, utilizing CRISPR-engineered TH-TdTomato reporter-expressing hiPSCs in both 2D monolayers and 3D mDA-astrocyte assembloids, addresses a critical gap in PD research by providing a scalable, human-relevant system to study disease mechanisms and identify therapeutic interventions. Our platform achieves high purity of SNpc-like mDA neurons expressing AGTR1, a receptor linked to PD-vulnerable neurons. The optimized protocols, including delayed dissociation and PEI + laminin coating, mitigate neuronal clumping, a common challenge in hiPSC cultures, ensuring precise automated quantification of mDA neuron survival via TH-TdTom fluorescence, segmentation, and automated counting. This model’s novelty lies in its ability to employ PD-relevant neurotoxic insults (CBE and Rotenone) to achieve robust and quantifiable human mDA neuronal loss with oxidative stress and accumulation of toxic forms of a-synuclein, enabling the identification of novel neuroprotective compounds and offering a translatable platform for therapeutic discovery in PD.

PD patient iPSC-derived mDA neurons have also been used to model PD in a dish [35–40]. Mitochondrial dysfunction, oxidative stress, and α-synuclein accumulation have been observed in these models, making them valuable to study pathogenic mechanisms. However, no or little DA neuronal loss is observed in these models, necessitating the use of neurotoxins to precipitate neurodegeneration phenotypes.

### Protection of human mDA neurons via AGTR1 inhibition

A key strength of our platform is its suitability for testing small molecules in a 96-well format, particularly for repurposing clinically approved drugs to mitigate neurodegeneration. Building on our previous findings in a zebrafish model [1,53], we identified AGTR1 inhibitors as potent neuroprotective agents in human mDA neurons against CBE-induced neurotoxicity, with candesartan showing near-complete protection at high doses. The high doses needed for neuroprotection in culture reflect that sartans are highly protein-bound [42]. Different levels of neuroprotection exerted by different sartans are likely due to their differential physicochemical properties, such as cell permeability and stability. Neuroprotective effects of sartans in human mDA neurons are further validated by genetic knockdown of AGTR1 via CRISPRi.

Compared to CBE, rotenone induced more severe mDA neurodegeneration, which was not effectively protected by AGTR1 inhibition. This observation suggests that neuroprotective strategies such as AGTR1 inhibition are best combined with the development of early disease diagnosis and detection technologies such as the α-synuclein seed amplification assay [54].

### Human-specific features of the RAAS pathway in the nigrostriatal system

Our analyses of the whole-brain single-nuclei transcriptomic data from humans and mice [22,23] revealed critical species-specific differences in RAAS pathway expression, underscoring the limitations of mouse models and the necessity of human-based systems for reliable neuroprotective strategies. In humans, AGTR1 is selectively expressed in PD-vulnerable TH⁺SOX6⁺CALB1⁻ SNpc neurons, whereas the equivalent mouse population lacks AGTR1a/b and expresses AGTR2, which is neuroprotective [34]. This divergence explains why AGTR1-mediated mechanisms, such as oxidative stress and neuroinflammation, are poorly recapitulated in mice, as AGTR2’s opposing role may mask AGTR1’s pro-degenerative effects. Additionally, the source of Renin in the brain has been subject to controversy. While mouse brain cells show little to no Renin expression [23], human Renin expression is found in the Choroid plexus and MSN neurons. Human SNpc astrocytes express AGT, the precursor to AGTR1’s ligand, and our human mDA platform, incorporating neuron-glia interactions, mirrors these human-specific RAAS dynamics, particularly the AGTR1-AGT axis in the nigrostriatal pathway. These findings highlight the need for human *in vitro* models to identify neuroprotective strategies that accurately reflect PD pathogenesis and ensure therapeutic relevance.

### The link between AGTR1 and α-synuclein in human mDA neurons

Our transcriptomic analyses of hiPSC-derived mDA neurons upon AGTR1 inhibition have revealed key biological processes impacted by AGTR1, including the level of α-synuclein. The transcripts of α-synuclein, β-synuclein, and SNCAIP were all significantly reduced by AGTR1 inhibitors, suggesting that AGTR1 functions to promote their transcription or transcript stability. In addition to transcriptional changes, AGTR1 inhibition significantly decreased phosphorylated α-synuclein at serine 129 (p129α-Syn). The p129α-Syn is a hallmark of pathological α-synuclein aggregates, such as Lewy bodies and Lewy neurites, observed in PD, dementia with Lewy bodies (DLB), and multiple system atrophy (MSA). Approximately 90% of α-synuclein in Lewy bodies is phosphorylated at Ser129, compared to only ∼4% in soluble forms, suggesting a strong association with disease pathology. Phosphorylation at Ser129 is thought to promote α-synuclein misfolding and aggregation, contributing to the formation of toxic oligomers and fibrils [50]. It is possible that AGTR1 inhibition primarily decreased synuclein transcripts, and such a reduction in synuclein expression led to reduced p129α-Syn. Alternatively, AGTR1 inhibition primarily decreased the level of p129α-Syn, through decreasing the activity of the kinases that carry out this phosphorylation event, and the reduction of p129α-Syn exerts a feedback regulation on the level of α-synuclein transcripts.

Given the high expression of α-synuclein in neurons, especially at presynaptic terminals, and its physiological role in regulating synaptic transmission, synaptic plasticity, and learning and memory [55–57], it is tempting to speculate that the AGTR1-αSyn pathway defined in this study may play a physiological role in regulating normal neuronal function, and that deregulation of this pathway under neurotoxic insults, chronic stress, and during aging may contribute to neurodegeneration and PD pathogenesis.

### Therapeutic outlook of AGTR1 inhibition for treating PD and other synucleinopathies

AGTR1 inhibitors are of wide clinical use to treat hypertensive patients, some of whom also have PD. Retrospective clinical studies, including ours, have yielded mixed results, reflecting individual variability and gaps in understanding the brain RAAS signaling. Whereas some studies suggest that known AGTR1 inhibitors reduce PD risk or slow progression [53,58,59], others report no significant benefits [60,61].

A challenge to translating AGTR1 inhibitors for PD is their limited blood-brain barrier (BBB) penetration [42], which may underlie the inconsistent clinical outcomes. While candesartan exhibits better BBB permeability than other sartans, its brain penetration remains suboptimal, as evidenced in a recent clinical trial [59]. Developing novel AGTR1 inhibitors with enhanced BBB penetrance is essential to maximize their potential therapeutic efficacy. Structure-activity relationship studies and nanoparticle-based delivery systems could improve CNS targeting, ensuring sufficient drug concentrations in the SNpc to modulate AGTR1 signaling effectively. With its high-throughput screening capabilities, our platform is ideally suited to test these next-generation inhibitors, accelerating their development for PD.

Our findings also provide a platform and support the rationale for combining AGTR1 inhibitors with additional pathway modulators to achieve synergistic neuroprotection. The platform we have established—including live TH-reporter imaging, RNA-seq pipelines, and scalable 3D assembloids—is well-suited for combinatorial candidate studies. For instance, combining sartans with other ROS-modulating agents, as suggested by the upregulation of *PRDX1* and *G6PD* upon AGTR1 inhibition in our transcriptomic data, could amplify antioxidant defenses, addressing oxidative stress, a central driver of PD pathology [34]. A recent study shows that α-Syn binds to and anchors G6PD to synaptic vesicles to ensure proper DA release. Loss of G6PD function can lead to PD phenotypes, and restoration of G6PD activity rescues DA signaling [45]. Our platform is also well-suited for next-generation functional genomics approaches. Given the high purity in mDA neurons we obtain and the possibility to sort pure mDA neuron populations based on TH-TdTom fluorescence, our model is well-suited for CRISPRi/a screens [62,63] that have yet to be performed on mDA neurons. In future studies, druggable-genome CRISPRi/a libraries can be leveraged to uncover additional modifiers of mDA neuron survival and identify therapeutic strategies that converge on lysosomal, inflammatory, and synuclein-regulating pathways, based on previous successful CRISPRi/a screens using hiPSC-derived neurons.

## LIST OF ABBREVIATIONS

PD: Parkinson’s Disease
mDA: Midbrain Dopaminergic
RAAS: Renin-Angiotensin-Aldosterone System
AGTR1: Angiotensin Receptor Type 1
hiPSC: Human Induced Pluripotent Stem Cell
TH: Tyrosine Hydroxylase
SNCA: Synuclein Alpha
SNCB: Synuclein Beta
SNCAIP: Synuclein Alpha Interacting Protein
p129α-Syn: Phosphorylated α-Synuclein at Serine 129
SNpc: Substantia Nigra Pars Compacta
GPCR: G-Protein Coupled Receptor
EB: Embryoid Body
GFR: Growth Factor Reduced
ROCK: Rho-Associated Protein Kinase
GDNF: Glial Cell Line-Derived Neurotrophic Factor
BDNF: Brain-Derived Neurotrophic Factor
TGFβ3: Transforming Growth Factor Beta 3
CNTF: Ciliary Neurotrophic Factor
FBS: Fetal Bovine Serum
CBE: Conduritol B-Epoxide
GBA1: Glucocerebrosidase
Rot: Rotenone
CALM: Center for Advanced Light Microscopy
FCS: Fluorescence Cytometry Software
PFA: Paraformaldehyde
DAPI: 4’,6-Diamidino-2-Phenylindole
RNA-seq: RNA Sequencing
snRNA-seq: Single-Nuclei RNA Sequencing
scRNA-seq: Single-Cell RNA Sequencing
DEGs: Differentially Expressed Genes
ROS: Reactive Oxygen Species
PRDX1: Peroxiredoxin-1
G6PD: Glucose-6-Phosphate Dehydrogenase
BBB: Blood-Brain Barrier
MSNs: Medium Spiny Neurons
PEI: Polyethylenimine
Lam: Laminin
PDL: Poly-D-Lysine
CRISPRi: CRISPR Interference
CRISPRa: CRISPR Activation
DMSO: Dimethyl Sulfoxide
α-Syn: Alpha-Synuclein
DLB: Dementia with Lewy Bodies
MSA: Multiple System Atrophy
FACS: Fluorescence-Activated Cell Sorting

## DECLARATIONs

### -Ethics approval and consent to participate

Not applicable.

### -Consent for publication

Not applicable. No individual person’s data or identifying information is included in this manuscript.

### -Availability of data and material

The datasets generated and/or analyzed during the current study are presented in the manuscript.

### Author’s contributions

S.G. and M.D. were responsible for project oversight and design. M.D. conducted experimental work and analyses with the assistance of V.M. for cell culture, Y.L., X.Y., and Y.S. for AGTR1-CRISPRi design and lentiviral particle production.

## Acknowledgements

We thank the Center for Advanced Light Microscopy (CALM) at UCSF, especially Caroline Mrejen, for her invaluable help and advice, and the members of the Guo lab for helpful discussions. We also thank Dr. Yin Shen and Dr. Bingwei Lu’s labs for sharing materials and expertise regarding CRISPRi technology and PD pathology. We acknowledge BIOSTATE for performing the RNA-seq analysis.

## FIGURES AND FIGURE LEGENDS

**FIGURE S1.**
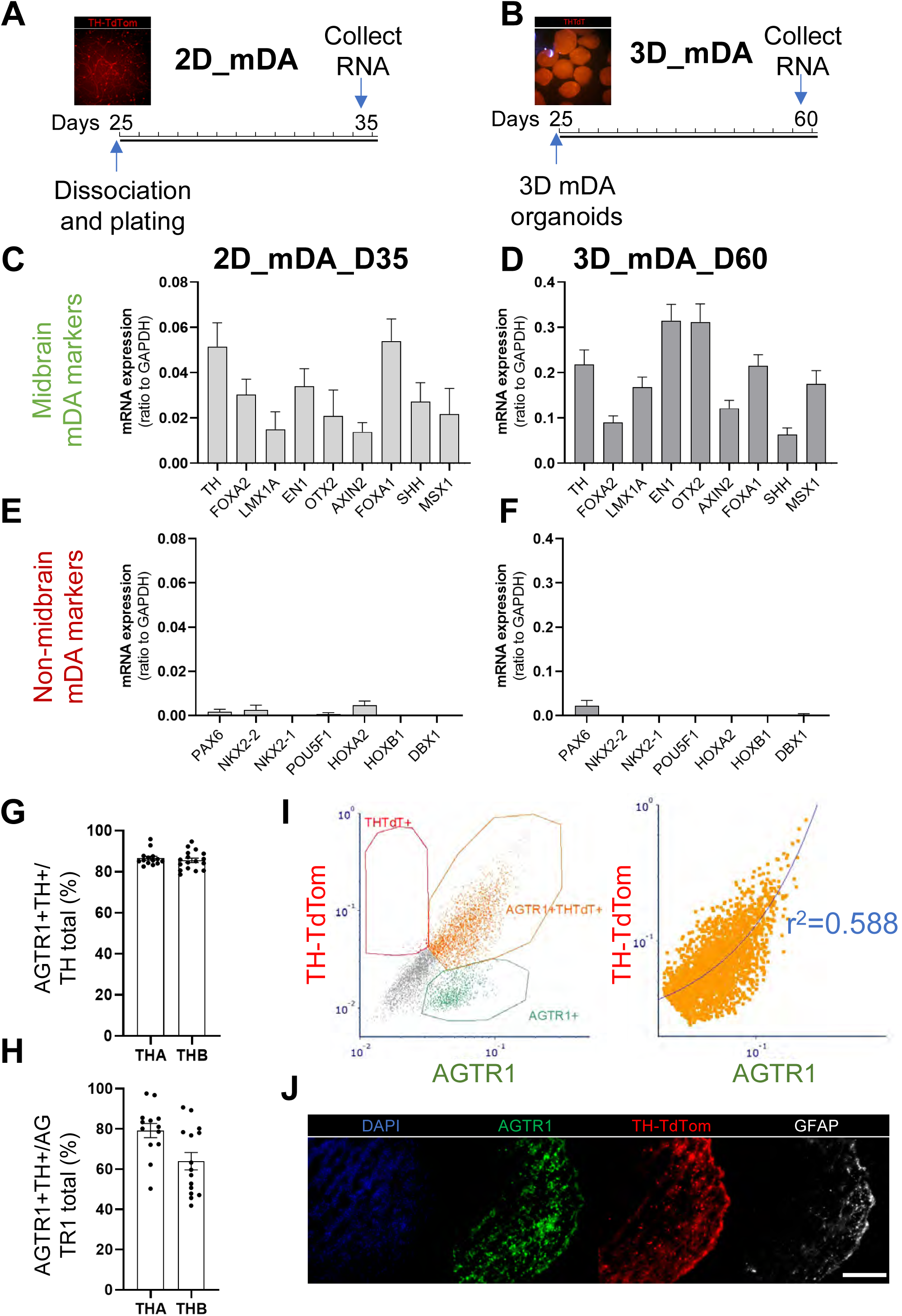
Gene expression analysis in 2D and 3D human mDA neuronal models. (**A, B**) Diagrams comparing 2D_mDA and 3D_mDA neuron models, indicating plating and RNA collection timelines. (**C, D**) mRNA expression in 2D vs 3D models for genes defining mDA identity. (**E, F**) mRNA expression in 2D vs. 3D models for genes expected to show low/no expression in mDA neurons. (**G**) Quantification of AGTR1+TH-TdTom+ cells as a proportion of total TH-TdTom+ cells in the mDA organoid model (D40). Two sub-lines (A and B) are shown. (**H**) Quantification of AGTR1+TH-TdTom+ cells as a proportion of total AGTR1+ cells in the mDA organoid model (D40). Two sub-lines (A and B) are shown. (**I**) A correlation plot of TH-TdTom vs. AGTR1 expression in mDA-mAstro assembloids, illustrating co-expression patterns. (**J**) A neuron-prominent region of an mDA-mAstro assembloid showing AGTR1 and TH-TdTom expression alongside GFAP+ glial infiltration. Scale bar: 200 μm (J).

**FIGURE S2.**
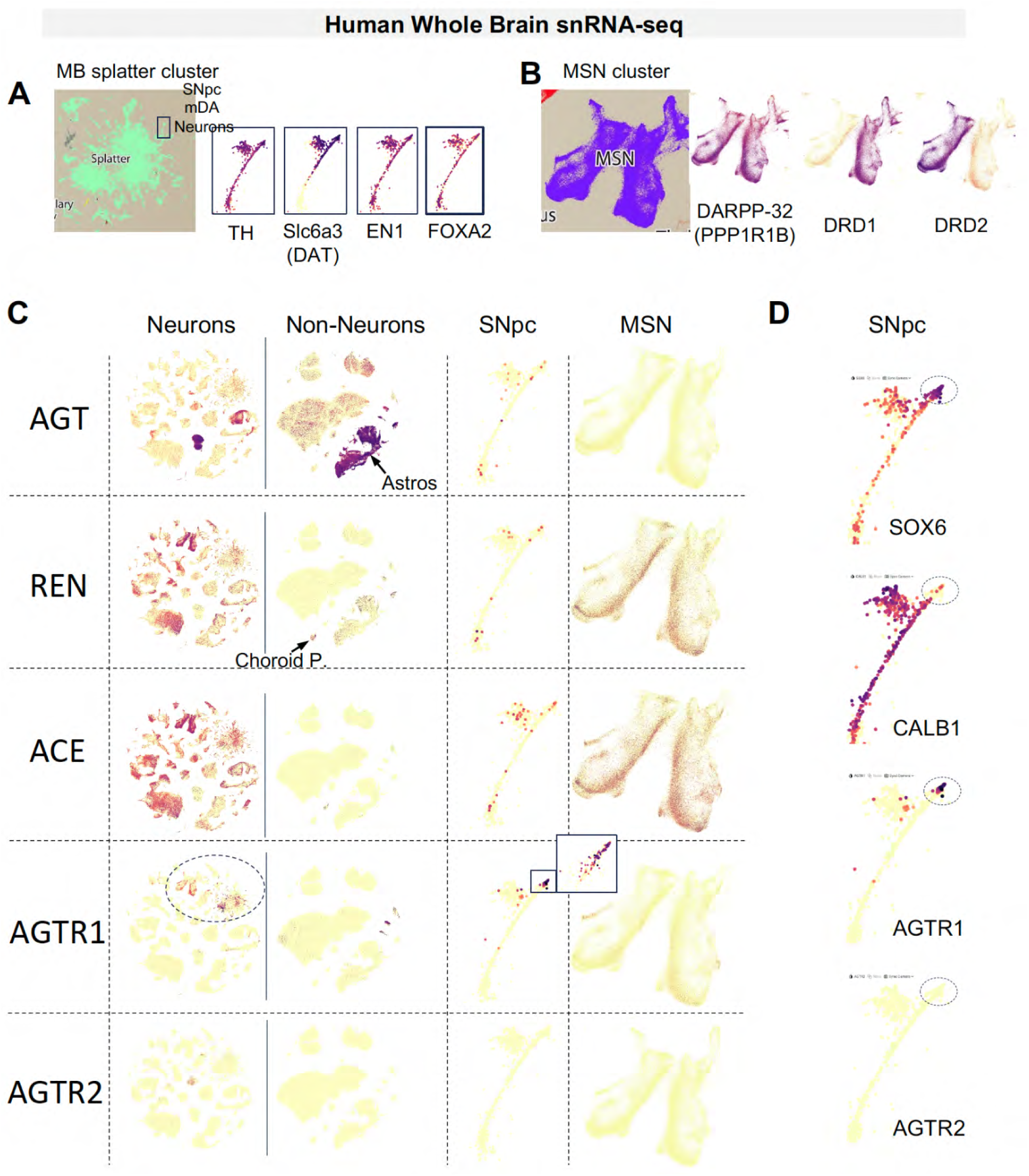
RAAS Pathway Component Expression in the human nigrostriatal pathway from the whole-brain snRNA-seq Database. (**A, B**) Images of substantia nigra pars compacta (SNpc) and Medium Spiny Neuron (MSN) clusters from the human whole brain high-resolution snRNAseq dataset identified by specific markers: SNpc mDA neurons express TH, DAT/SLC6A3, FOXA2, and EN1; MSNs express DARPP-32 (PPP1R1B), DRD1, and DRD2. (**C**) Images show AGT expression in astrocytes and AGTR1 expression in a subset of SNpc mDA neurons. Interestingly, striatal MSNs express both REN and ACE and low AGTR1. (**D**) Focused images of human SNpc mDA neurons identified in (A) show expression of AGTR1 in TH+SOX6+CALB1-neurons (dotted circle) with no expression of AGTR2.

**FIGURE S3.**
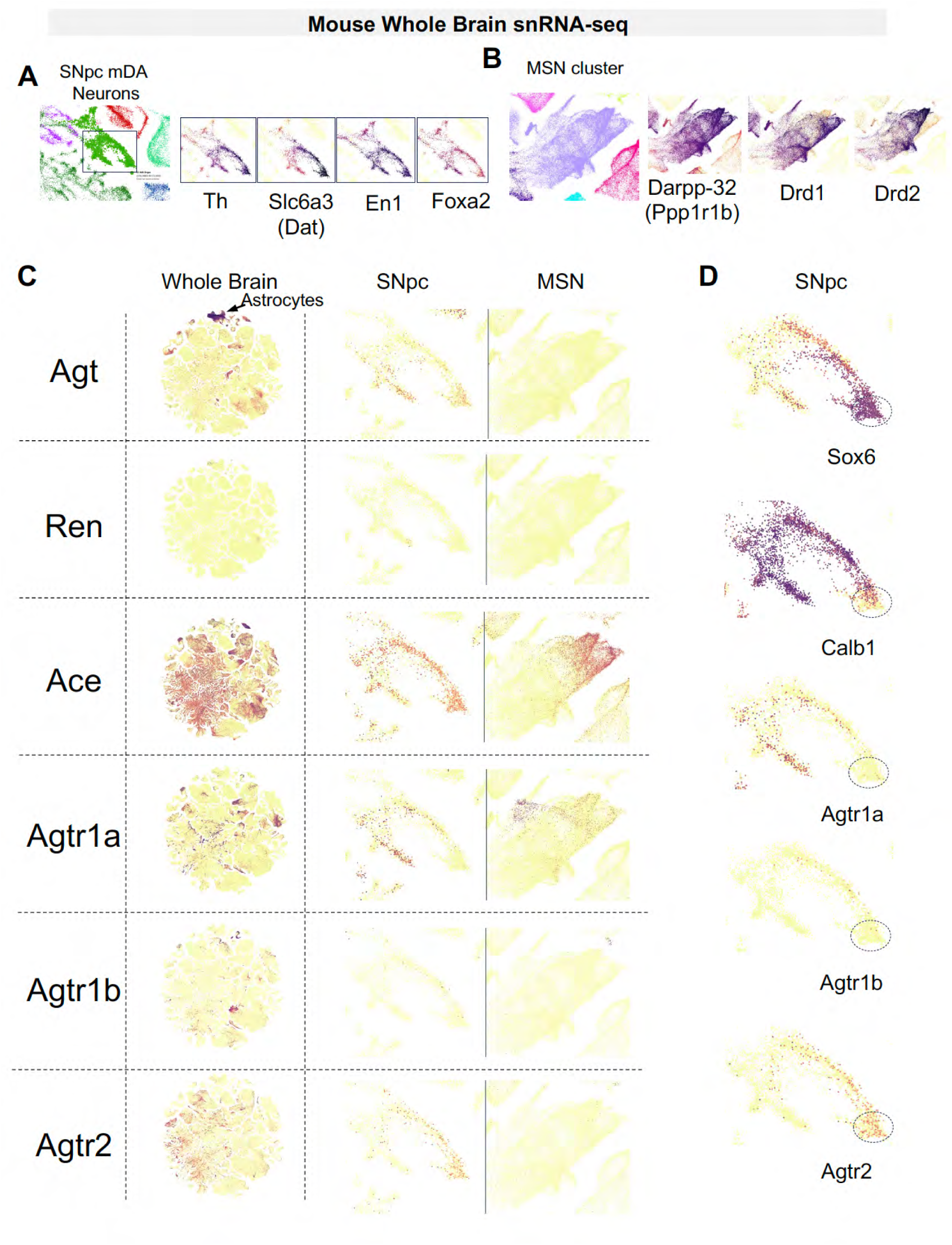
RAAS Pathway Component Expression in the mouse nigrostriatal pathway from the whole-brain snRNA-seq Database. (**A, B**) Images of substantia nigra pars compacta (SNpc) and Medium Spiny Neuron (MSN) clusters in the mouse whole brain snRNAseq database identified by specific markers: SNpc mDA neurons express Th, Dat/Slc6a3, Foxa2, and En1; MSNs express Darpp-32 (Ppp1r1b), Drd1, and Drd2. (**C**) Images show Agt expression in astrocytes. Unlike the human brain, Ren shows minimal to no expression throughout the brain. Ace is broadly expressed, with specific enrichment in striatal MSNs. Agtr1a and Agtr1b are expressed across various neuronal types, more broadly than in humans, with Agtr1a detected in SNpc and MSN neurons, but minimal Agtr1b in these regions. Unlike humans, Agtr2 is broadly expressed in the brain, including SNpc neurons. (**D**) Focused images of mouse SNpc mDA neurons from (A) show Agtr2 expression in Th+Sox6+Calb1-neurons (dotted circle), but no Agtr1a or Agtr1b expression, contrasting with human SNpc neurons, highlighting species-specific differences in the brain RAAS pathway.

**FIGURE S4.**
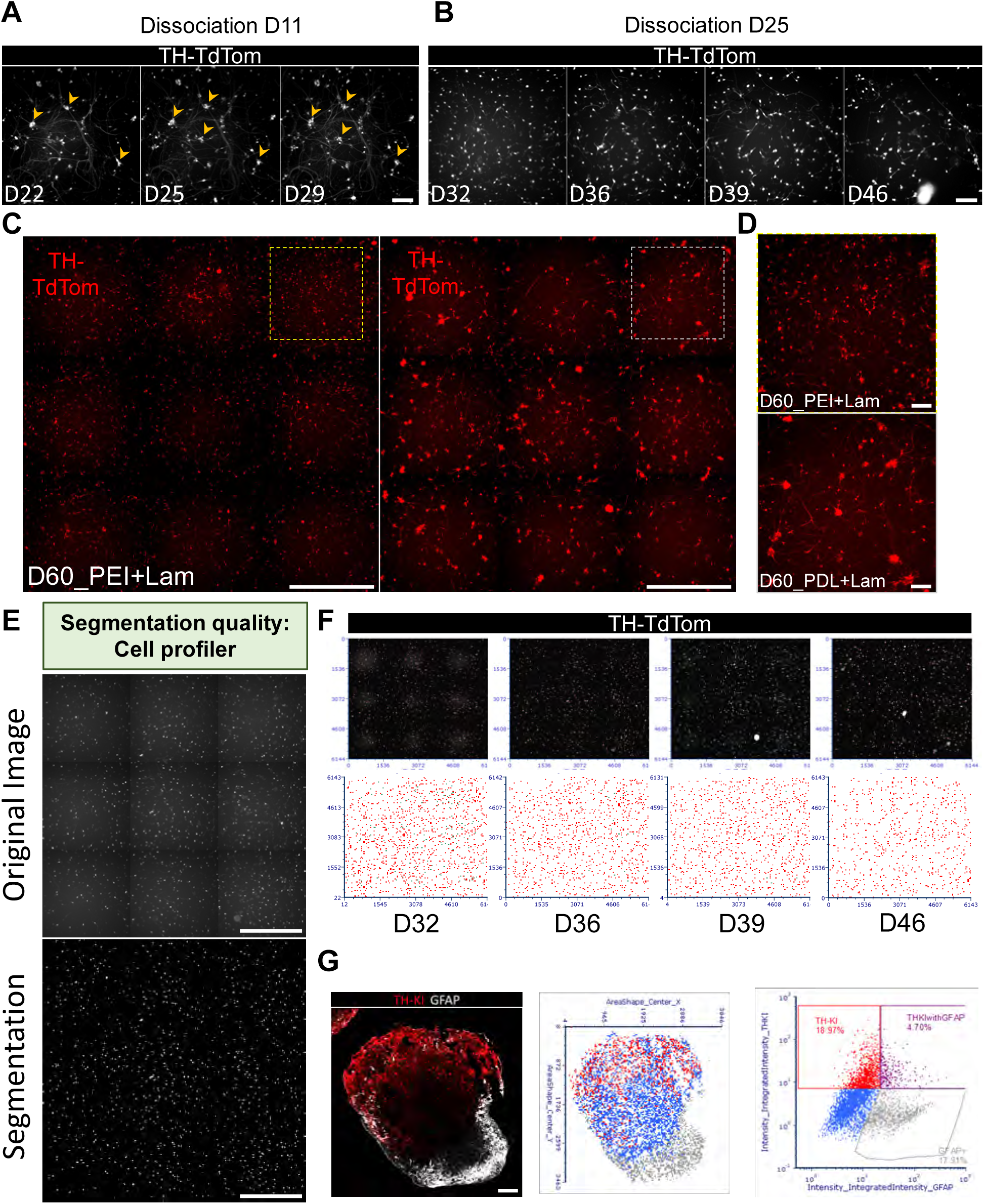
High content 2D and 3D human mDA neuronal culture, imaging, and quantification. (**A, B**) Neuronal clumping was prevented by dissociation at D25 compared to D11. (**C, D**) Examples of differences in clumping between PEI+Lam and PDL+Lam coating; only PEI+Lam coating allows long-term avoidance of neuron clumping, a prerequisite for reliable neuronal counting. (**E**) Segmentation quality of a whole well using a customized Cell Profiler 4.0 pipeline. (**F**) Survival experiment example for an entire well, displaying TH-TdTom+ images with corresponding FCS Express 7.0 image cytometry characterization (red dots indicate cells). (**G**) An example of counting on an mDA-mAstro assembloid section, using Cell Profiler 4.0 + FCS Express 7.0 image cytometry strategy. Scale bars: 1 mm (C, E); 100 μm (A, B, D, G).

**FIGURE S5.**
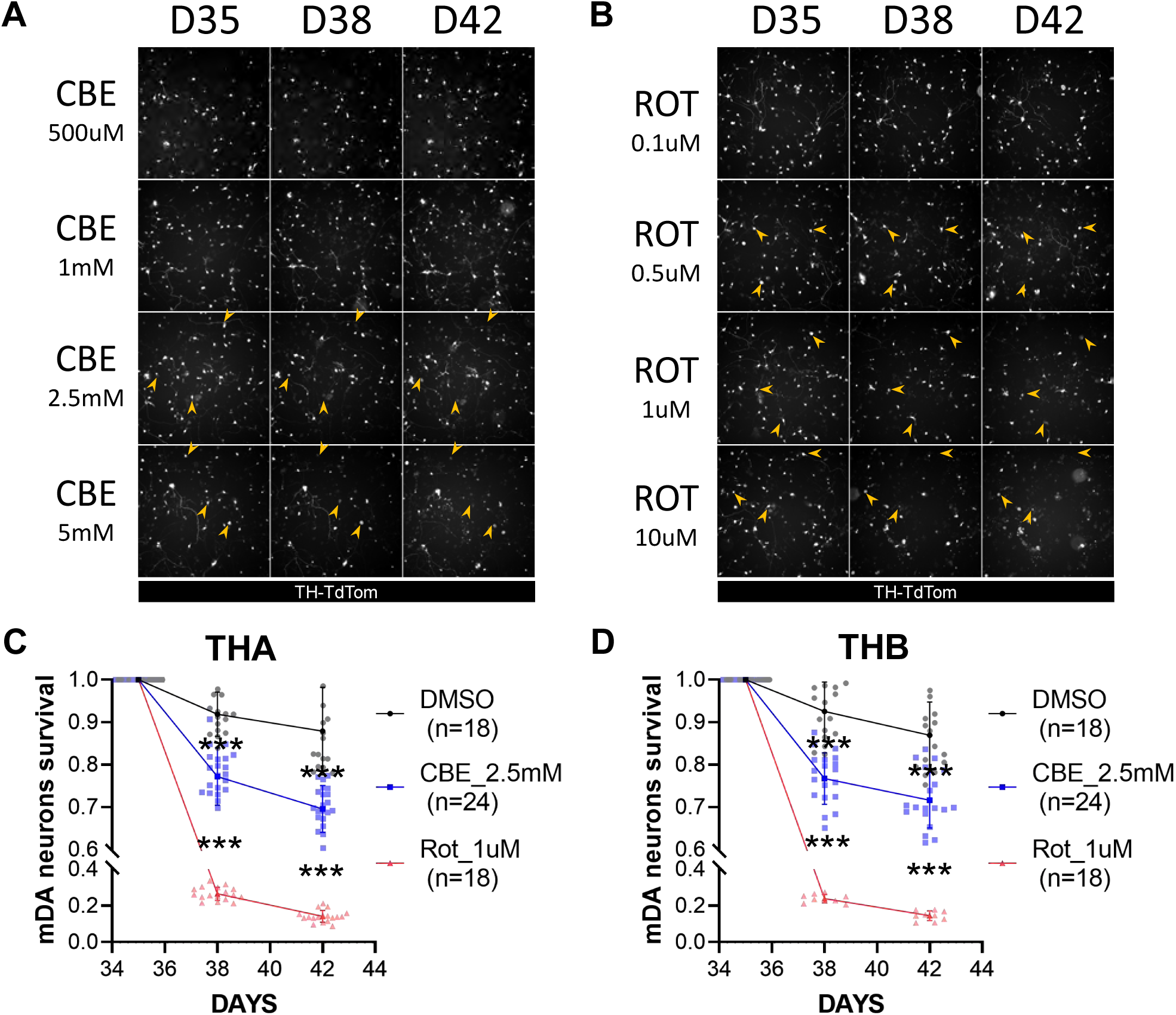
CBE- and Rotenone-induced human mDA neurodegeneration models in a dish. (**A, B**) To induce neurodegeneration, CBE [500μM-5mM] and Rotenone [0.1μM-10μM] were used at different concentrations on mDA neurons at D35 in culture to evaluate the impact on neuronal loss at D38 and D42. An example of cells dying at different concentrations of CBE and Rot is shown here with yellow arrowheads. (**C, D**) [CBE]_2.5mM or [Rot]_1μM treatment on two different clones of the TH-TdTom KOLF2.1 hiPSC line (THA and THB) shows the reproducibility and robustness of our chemically induced neurodegeneration models. *** p<0.001. Scale bars: 100 μm.

**FIGURE S6.**
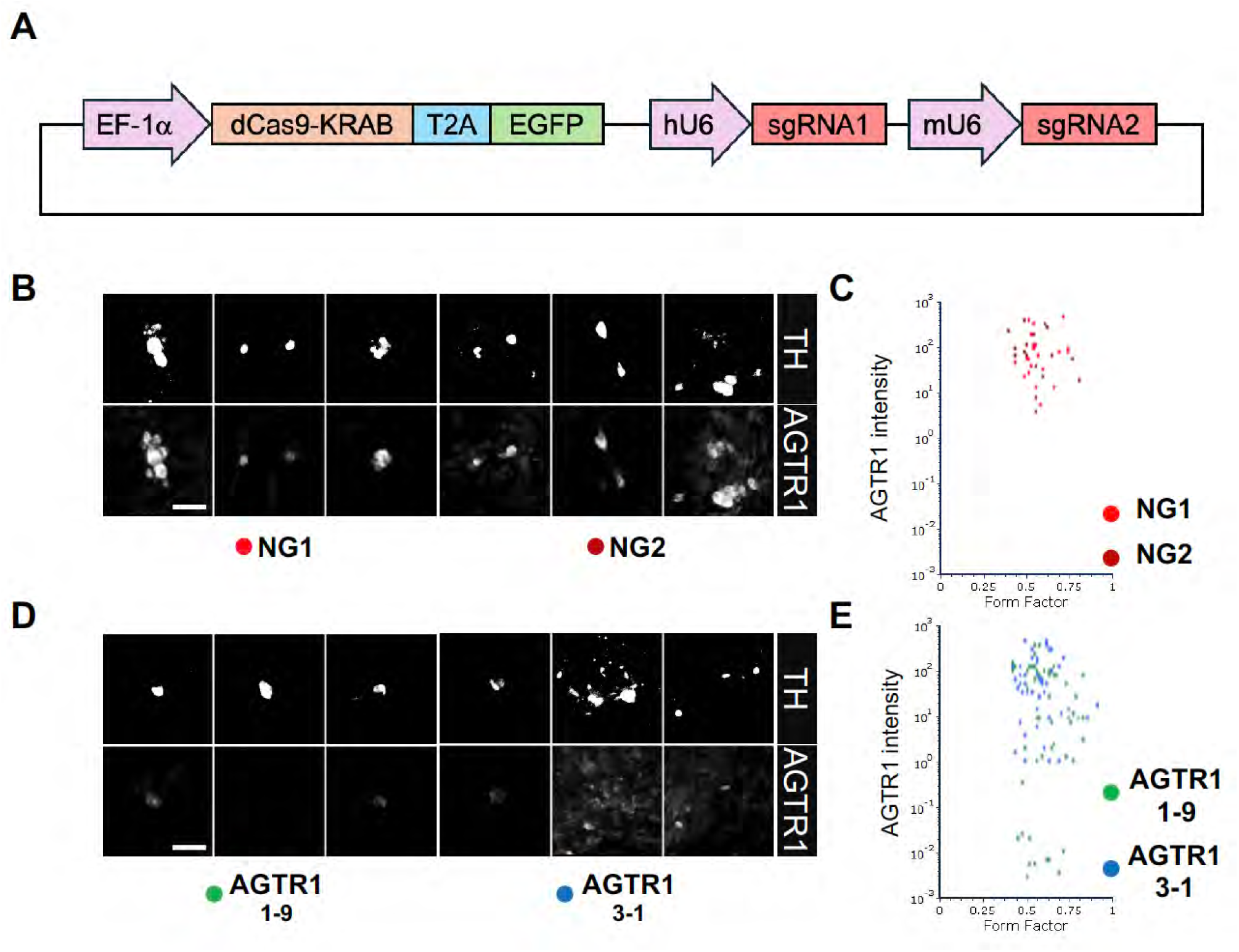
AGTR1-CRISPRi design and validation. (**A**) AGTR1-CRISPRi design strategy. (**B, C**) Example of AGTR1-expressing neurons using control NG1 and NG2 lentiviral particles, quantification in C. (**D, E**) Example of AGTR1-KD neurons using control AGTR1-9 and AGTR3-1 lentiviral particles, quantification in E, showing partial AGTR1 knockdown. Scale bars: 30 μm.

**FIGURE S7.**
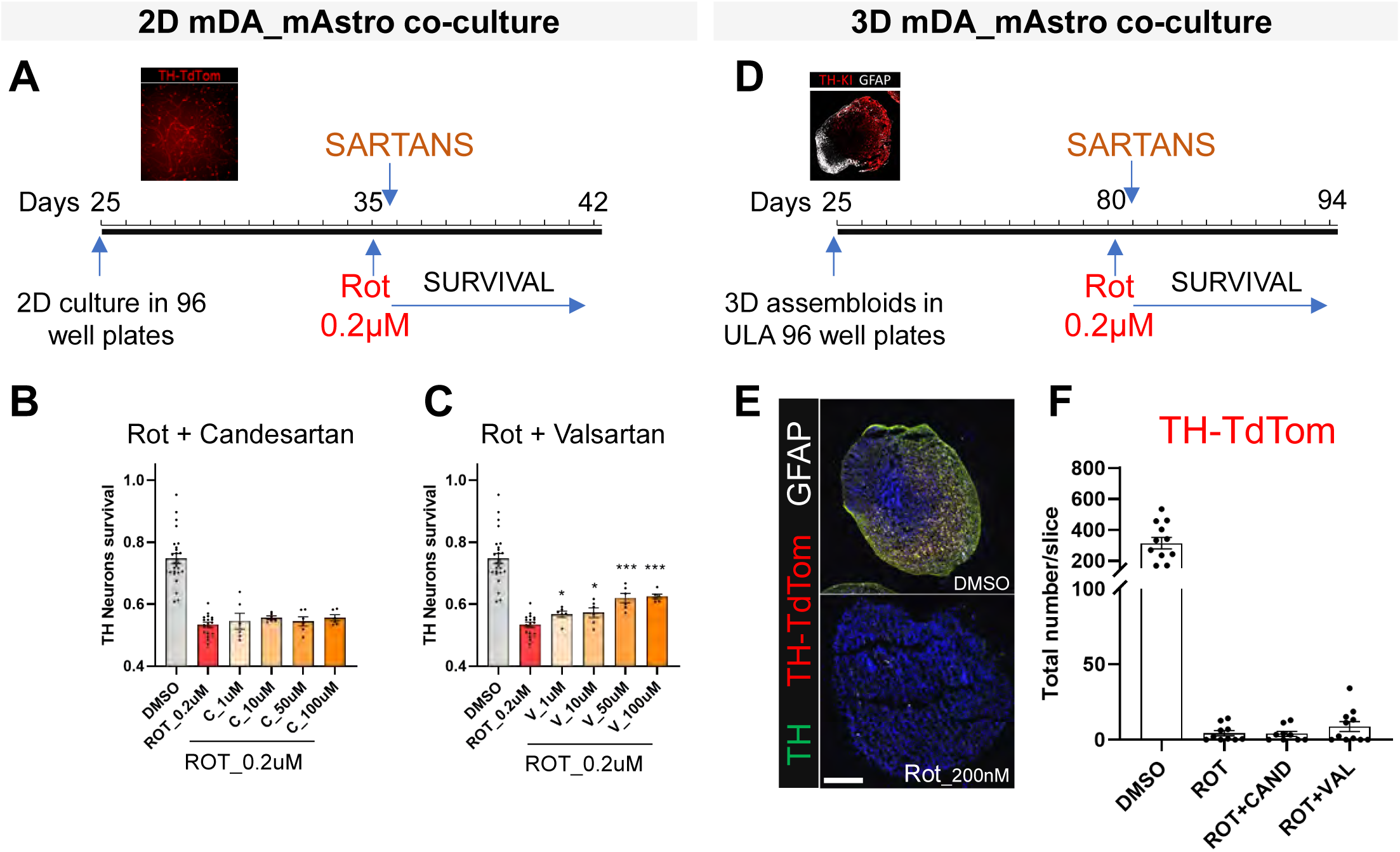
AGTR1 inhibitors show limited neuroprotection against the severe mDA neuron degeneration induced by rotenone. (**A, D**) Pipeline of survival experiments in 2D mDA-mAstro co-culture and 3D mDA-mAstro assembloids. Rot treatment was followed by AGTR1 inhibitors (sartans). (**B, C**) Candesartan did not show neuroprotection against low Rot dose [0.2 μM] while high concentrations of Valsartan showed partial neuroprotection. (**E**) Even a low Rot dose [0.2 μM] shows high neurotoxicity in the 3D mDA-mAstro assembloid model. (**F**) Quantification of TH-TdTom+ neurons on mature mDA-mAstro assembloids (D80) shows that neither Candesartan nor Valsartan can counteract Rot’s high toxicity. Scale bars: 200 μm.

**FIGURE S8.**
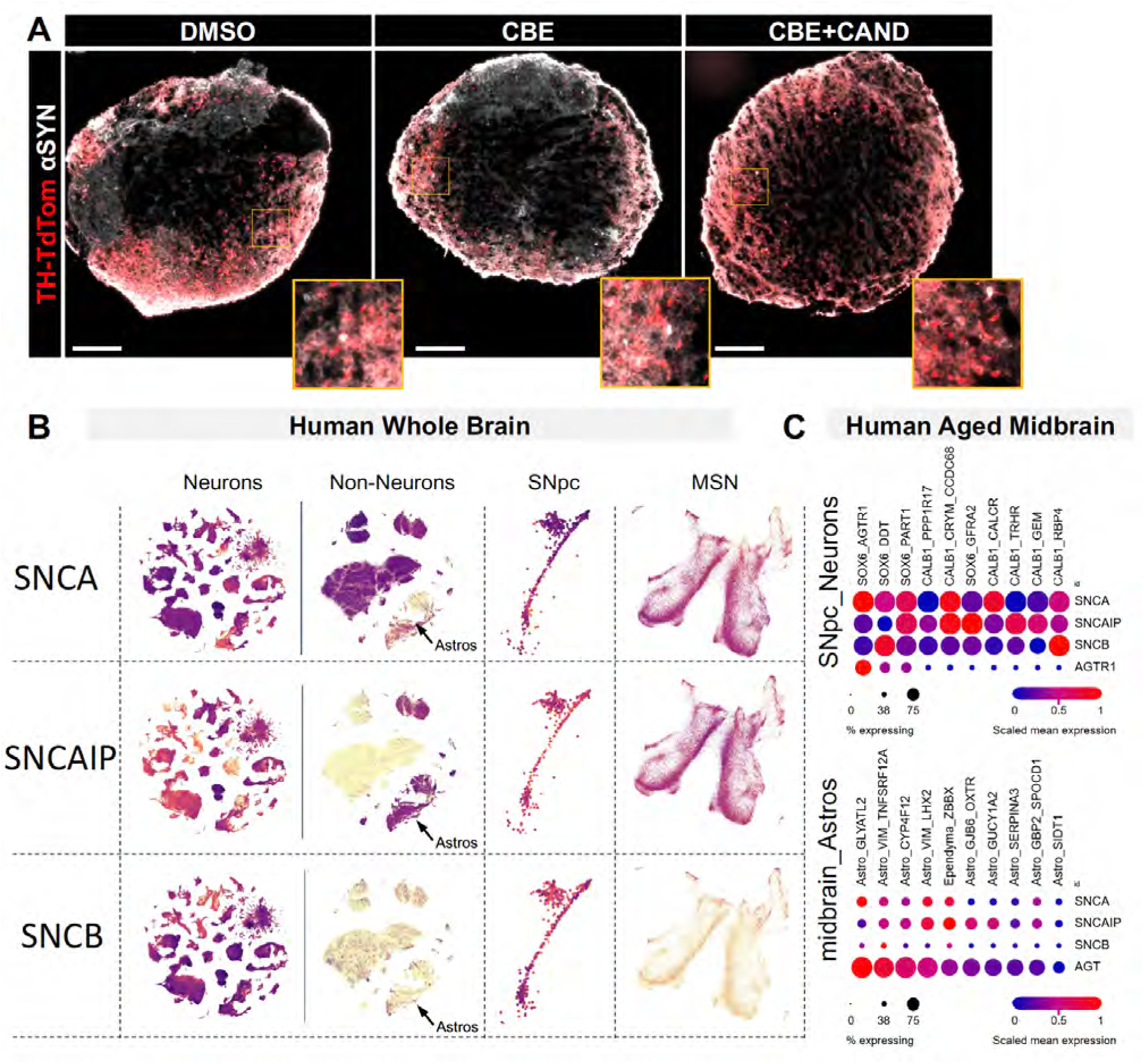
αSyn immunofluorescent labeling on 3D mDA-mAstro assembloids and the expression of synucleins in the human snRNA-seq database. (**A**) Immunofluorescent labeling of α-Synuclein on D74 mDA-mAstro assembloids from Fig. 6D. (**B, C**) Images from the human whole-brain snRNA-seq database and the Human Aged Midbrain database show high expression of SNCA, SNCAIP, and SNCB in neuronal populations and SNpc. However, only SNCA and SNCAIP, and not SNCB, show high expression in Astrocytes and MSN neurons. Scale bars: 200 μm.

**FIGURE S9.**
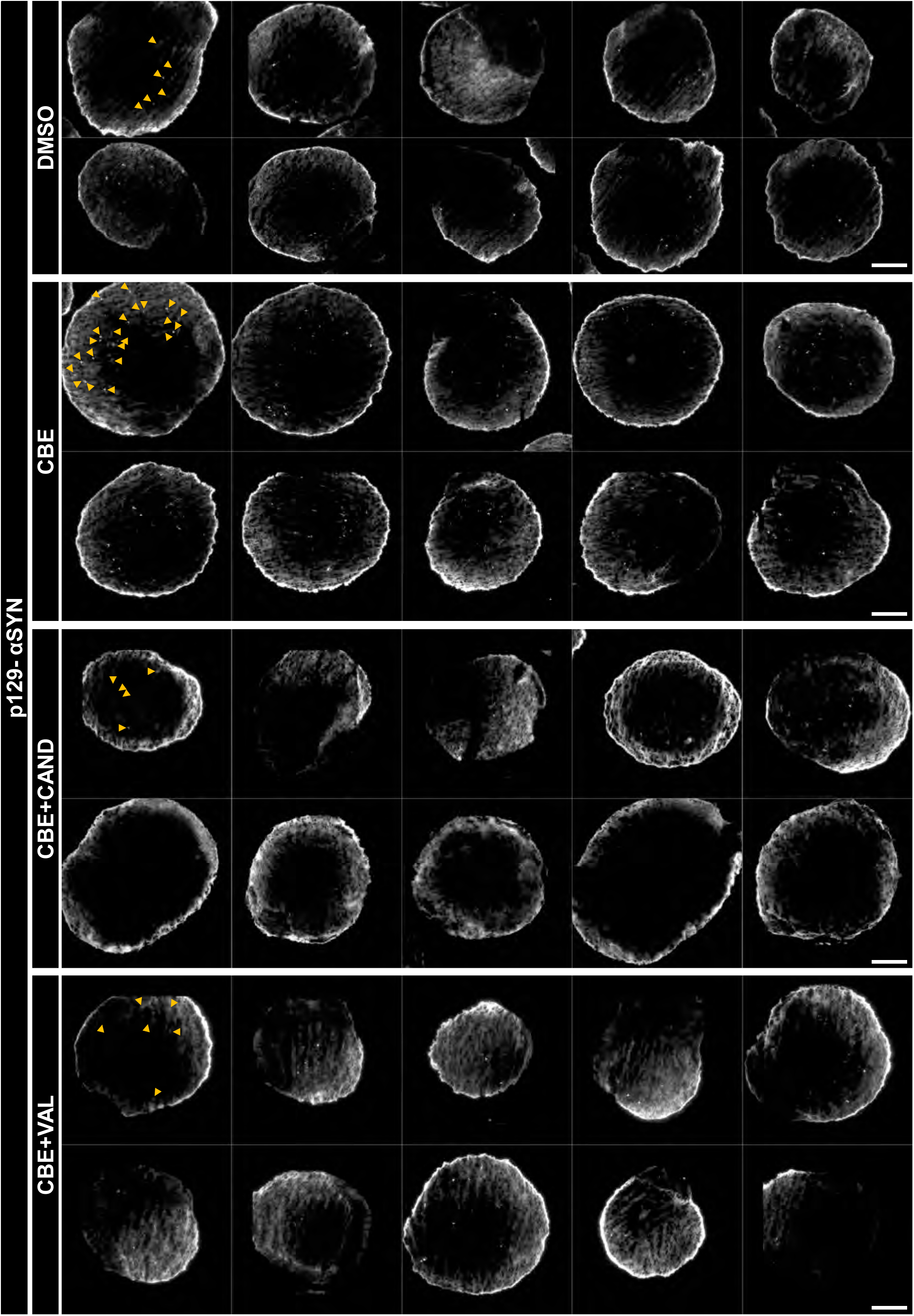
Composites of 3D mDA-mAstro assembloids immunofluorescently labeled with p129aSyn. Examples of p129-αSyn staining on 10 random 3D mDA-mAstro assembloid slices for each condition, as represented in Fig. 6F, show a consistent increase of p129-αSyn positive cells in the CBE condition, and a reproducible decrease in SARTAN-treated conditions. Examples of p129-αSyn positive cells are shown with yellow arrow heads on the top left image for each condition. Scale bars: 200 μm.

